# Evasion of Hepatic ILC1 Immunosurveillance Drives Breast Cancer Liver Metastasis

**DOI:** 10.64898/2026.10.26.759951

**Authors:** Peng Li, Jun Li, Andrew E. Cornish, Jing Zhang, Xian Zhang, Dorukhan Bahceci, Zijian Xu, Liangliang Ji, Si-Kun Guo, Sophia Brosnan, Briana G. Nixon, Juan Yan, Ping Chi, Fresia Pareja, Pedram Razavi, Christopher Fong, Adam Price, Allison Richards, Ming O. Li

## Abstract

Metastasis, the dissemination of cancer cells to distant organs, is an inefficient process^1^, yet remains the leading cause of cancer-related mortality^2^. The cellular barriers and evasion mechanisms governing organ-specific susceptibility remain incompletely understood^3–5^, hindering the development of effective anti-metastatic therapies^6^. Here, we show that liver-resident type 1 innate lymphoid cells (ILC1s) constitute an essential immunosurveillance barrier against breast cancer liver metastasis that can be bypassed through either cancer cell-intrinsic or host-dependent mechanisms. In a cohort of patients with multi-organ metastatic breast cancer, liver metastasis occurred later and was associated with poorer survival than lung metastasis, with elevated body mass index emerging as a clinical risk factor. This metabolic vulnerability was recapitulated in mice, where a high-fat diet induced hepatosteatosis and selectively enhanced breast cancer colonization of the liver. Mechanistically, liver and lung colonization were restrained by ILC1s and circulating natural killer (NK) cells, respectively. Following cancer cell challenge, NK cells expanded and became activated, clearing cancer cells through the lytic granule pathway and engagement of the death receptor DR5, whereas activated ILC1s selectively required high DR5 expression on cancer cells, supported by AP-2α (TFAP2A)-mediated transcriptional activation, for effective immunosurveillance. Heterozygous copy-number deletion at *TNFRSF10B* (encoding DR5) in human tumours reduced DR5 expression and predisposed to liver metastasis. Hepatosteatosis further compromised this barrier by reducing IL-15 expression in macrophages, thereby depleting hepatic ILC1s, effects reversible with glucagon-like peptide-1 (GLP-1) receptor agonists. Thus, hepatic ILC1s form a critical organ-specific barrier to breast cancer metastasis, vulnerable to both cancer cell-intrinsic DR5 downregulation and host metabolic dysfunction, the latter of which is pharmacologically correctible.

## Main Text

Along the hematogenous route of metastatic dissemination, the liver is the largest and one of the most accessible target organs, uniquely distinguished by its dual blood supply from the hepatic portal vein and hepatic artery, fenestrated sinusoidal endothelium, and low intravascular pressure, features thought to facilitate the hepatic seeding of circulating cancer cells^7^. Yet whereas portal drainage drives the high incidence of liver metastases from gastrointestinal cancers, systemically disseminated cancer cells display variable propensities to colonize the liver relative to other organs^3^. Breast cancer, which frequently metastasizes to multiple distant sites, exemplifies this heterogeneity^8^. How cancer cell-intrinsic determinants and tissue-specific variables interact to dictate metastatic organotropism, as originally conceptualized in the “seed and soil” hypothesis^9^, remains incompletely defined.

### Dietary modeling of delayed liver metastasis

To investigate evolving trajectories of organotropic metastasis, we analyzed the temporal patterns of metastatic occurrence in an MSK-Integrated Mutation Profiling of Actionable Cancer Targets (MSK-IMPACT) Clinical Sequencing Cohort, focusing on breast cancers with multi-organ dissemination. Lung metastasis occurred at comparable frequencies across early and late time points, whereas bone metastasis was enriched among early events and liver and brain/central nervous system (CNS) metastases among later events (Fig. 1a, Extended Data Fig. 1a). Consistent with this pattern, liver and CNS/brain metastases were associated with poorer patient survival (Fig. 1b). The delayed emergence of liver metastasis is particularly notable, given that, unlike the brain, which is protected by the blood-brain barrier, the liver is not anatomically shielded from circulating cancer cells^3^. These findings suggest that breast cancer metastasis encounters distinct suppressive mechanisms within the hepatic niche that are progressively weakened or circumvented over time. Notably, a body mass index (BMI) value above the ethnic-inclusive World Health Organization optimal range^10^ of 23 kg/m^2^ emerged as an epidemiological risk factor for liver metastasis in our cohort (Fig. 1c, Extended Data Fig. 1b). Biomarker-based subtype analysis further showed that the BMI-associated liver metastasis risk was strongest in triple-negative (TN) breast cancer and less pronounced in hormone-receptor-positive (HR+) or HER2+ subtypes (Extended Data Fig. 1c).

**Fig. 1.**
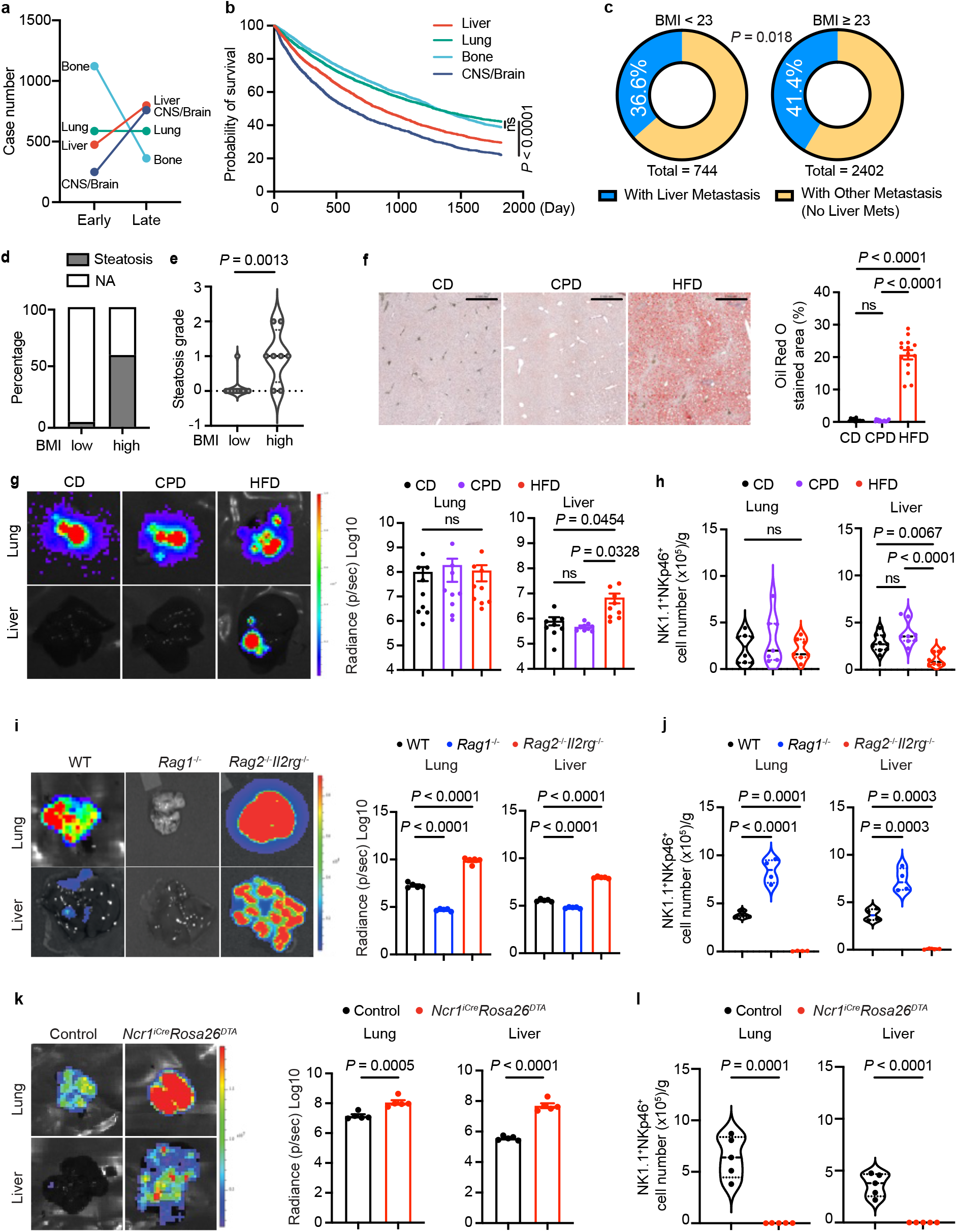
Depletion of group 1 innate lymphoid cells in high-fat diet-fed mice models the elevated body mass index-associated delayed onset of breast cancer liver metastasis. **a**, Metastatic onset at bone, lung, liver, and central nervous system (CNS)/brain sites in breast cancer patients. Early and late cases are defined as within one year or more than one year after the initial detection of metastasis, respectively. **b**, Breast cancer patient survival following organ-specific metastasis to bone (n = 4,096), liver (n = 3,102), lung (n = 3,063), and CNS/brain (n = 2,148). Survival probabilities were estimated with the Kaplan–Meier method; curves were compared with the two-sided log-rank (Mantel–Cox) test. *P* < 0.0001 for liver versus lung, liver versus bone and liver versus CNS/brain. **c**, Association between pre-metastatic body mass index (BMI) and the likelihood of breast cancer metastasis to liver versus other organs. BMI was measured >90 days before the first metastatic diagnosis. The BMI range (23 kg/m^2^) was used as the cut-off. **d**, Percentage of patients diagnosed with hepatic steatosis via imaging among individuals with liver metastasis, stratified into high and low BMI groups (n = 25 per group). Steatosis classification was determined by documented imaging records (MRI, CT, or ultrasound) or a documented NASH diagnosis. NA indicates patients without available or documented imaging data. **e**, Steatosis grades determined by pathology assessment for patients with liver biopsy samples (high BMI, n = 8; low BMI, n = 15). **f**, Representative images of Oil Red O staining and hematoxylin and eosin (H&E) staining, along with Oil Red O quantification, of liver tissue from C57BL/6 mice. Mice were fed either a control chow diet (CD), a high-fat diet (HFD; 60% kcal fat), or a control purified diet (CPD; calorie-matched with 10% kcal fat). Oil Red O staining quantification is shown as mean ± s.e.m. Scale bar: 0.5 mm. **g**, **i**, **k**, Representative bioluminescence images and radiance quantification of lung and liver colonization following intravenous administration of E0771 breast cancer cells in C57BL/6 mice fed a CD, CPD, or HFD (**g**), in wild-type (WT), *Rag1^−/−^,* or *Rag2^−/−^Il2rg^−/−^*mice (**i**), and in *Ncr1^iCre^Rosa26^DTA^* and littermate control mice (**k**) (n = 5 per group). Data are representative of at least three independent experiments, each cohort is n = 5 per group, one representative image shown. Radiance quantification is shown as mean ± s.e.m. **h**, **j**, **l**, Quantification of NK1.1^+^NKp46^+^ group 1 innate lymphoid cells in lungs and livers of C57BL/6 mice fed a CD, CPD or HFD (**h**), WT, *Rag1^−/−^,* or *Rag2^−/−^Il2rg^−/−^*mice (**j**), and *Ncr1^iCre^Rosa26^DTA^* and littermate control mice (**l**) before E0771 cancer cell challenge. Lymphocyte numbers per gram of tissue are shown as mean ± s.e.m. Mice were analyzed without tumor challenge. *P* values were calculated using the log-rank (Mantel-Cox) test (**b**), chi-squared test (**c**), Mann-Whitney *U* test (**e**), Welch’s *t* test (**k-l**), and Ordinary one-way ANOVA test (**f-j**). ns: not significant (*P* > 0.05).

Consistent with the metabolic functions of the liver and prior evidence linking hepatosteatosis to breast cancer liver metastasis recurrence^11^, patients in the high BMI cohort exhibited both increased prevalence of hepatosteatosis and elevated disease severity compared with those with low BMI (Fig. 1d, e). To experimentally model this BMI-associated metabolic vulnerability, C57BL/6 mice were maintained on a high-fat diet (HFD; 60% kcal from fat) for 16 weeks, resulting in substantial increases in body weight compared with control chow diet (CD)-fed mice (Extended Data Fig. 1d). To assess breast cancer metastasis susceptibility, we used E0771.lmb, a breast cancer cell line derived from a spontaneous tumour in C57BL/6 mice that exhibits TN breast cancer characteristics^12^. E0771.lmb cells expressing a luciferase reporter were orthotopically transferred to the mammary fat pad of CD- or HFD-fed mice, followed by primary tumour resection and monitoring for metastasis progression. HFD-fed mice developed larger primary tumours and exhibited greater metastasis susceptibility, particularly in the liver (Extended Data Fig. 1e-h), supporting high BMI-associated liver metastasis in patients.

To further characterize the HFD effects, we also fed mice a calorie-matched purified diet with lower fat content (CPD; 10% kcal from fat) (Extended Data Fig. 2a). Compared with CD- or CPD-fed mice, HFD-fed mice showed selective increases in body and liver weights (Extended Data Fig. 2b, c), with pronounced hepatic lipid accumulation evident by Oil Red O staining (Fig. 1f). Serum aspartate aminotransferase (AST) and alanine aminotransferase (ALT) levels remained unchanged (Extended Data Fig. 2d); yet, a number of serum lipid species, characteristic of patients with hepatosteatosis^13,14^, were significantly increased in HFD-fed, but not CD- or CPD-fed mice (Extended Data Fig. 2e, Extended Data Table 1). To further define the diet-associated liver metastasis susceptibility and rule out the effects of differences in primary tumour growth observed in the fat pad model, we administered E0771.lmb cells through intravenous inoculation to mimic systemic hematogenous dissemination of breast cancer cells. Robust liver colonization occurred exclusively in HFD-fed, but not CD- or CPD-fed mice, without altering the more readily observed dissemination to the lung (Fig. 1g, Extended Data Fig. 3a). To validate these findings in an independent model, we used PY8119 cells (a breast cancer cell line derived from MMTV-PyMT mice) depleted of B2M to mitigate confounding by the immunogenicity of the PyMT viral oncoprotein in transfer experiments (Extended Data Fig. 3b). Intravenous inoculation of PY8119 *B2m^−/−^* cells recapitulated HFD-associated liver metastasis susceptibility as observed in E0771.lmb cells (Extended Data Fig. 3c-d). These findings demonstrate that HFD-induced hepatic steatosis defines a permissive niche for breast cancer liver metastasis, offering an environmental explanation for its delayed occurrence observed in the clinic.

### Hepatic ILC1 suppression of liver metastasis

In a multiomic study of metastatic human breast cancer, liver, but not lung, metastases exhibited diminished immune scores compared with primary tumours^15^, a difference potentially attributable to organ-specific set points distinguishing the liver from the lung^16^. In agreement with these observations, enumeration of immune cell populations revealed selective depletion of natural killer T (NKT) cells, γδ T cells, and NK1.1^+^NKp46^+^ group 1 innate lymphoid cells (ILCs) in the livers, but not lungs, of HFD-fed mice (Fig. 1h, Extended Data Fig. 4). To delineate the relative contributions of distinct lymphoid lineages to anti-metastatic immunity, we employed *Rag1^−/−^* mice lacking adaptive lymphocytes and *Rag2^−/−^Il2rg^−/−^* mice devoid of both adaptive and innate lymphocytes. Whereas E0771.lmb cells robustly colonized both the lung and liver of *Rag2^−/−^Il2rg^−/−^*mice, their metastatic potential was markedly diminished in *Rag1^−/−^*mice relative to wild-type controls (Fig. 1i). These opposing metastatic phenotypes were inversely correlated with group 1 ILC abundance (Fig. 1j, Extended Data Fig. 5a). Importantly, selective depletion of group 1 ILCs, either genetically in *Ncr1^iCre^Rosa26^DTA^* mice^17^ or through anti-NK1.1 antibody treatment, significantly increased metastatic tumour burden in both lung and liver (Fig. 1k, l; Extended Data Fig. 5b-e). While neither approach is entirely specific to group 1 ILCs, the concordant results from these orthogonal depletion methods support a critical role for these cells in metastasis immunosurveillance in the liver, and, to a lesser extent, in the more susceptible lung.

Group 1 ILCs comprise both conventional natural killer (NK) cells, which recirculate through the bloodstream, and type 1 innate lymphoid cells (ILC1s), which reside in tissues^18^. Whereas CD49a^−^CD49b^+^ NK cells predominated in the lung, both NK cells and CD49a^+^CD49b^−^ ILC1s populated the liver (Fig. 2a). Following E0771.lmb challenge, NK cells, but not ILC1s, expanded (Fig. 2a), accompanied by increased frequencies of proliferating Ki67^+^ cells (Fig. 2b). To delineate NK cell contributions to metastasis immunosurveillance, we used anti-asialo-GM1 antibodies, which efficiently depleted NK cells without substantially affecting ILC1s. NK cell depletion increased lung, but not liver, metastasis (Fig. 2c, d; Extended Data Fig. 6a, b). Thus, NK cells restrain breast cancer lung metastasis but are dispensable for liver metastasis control.

**Fig. 2.**
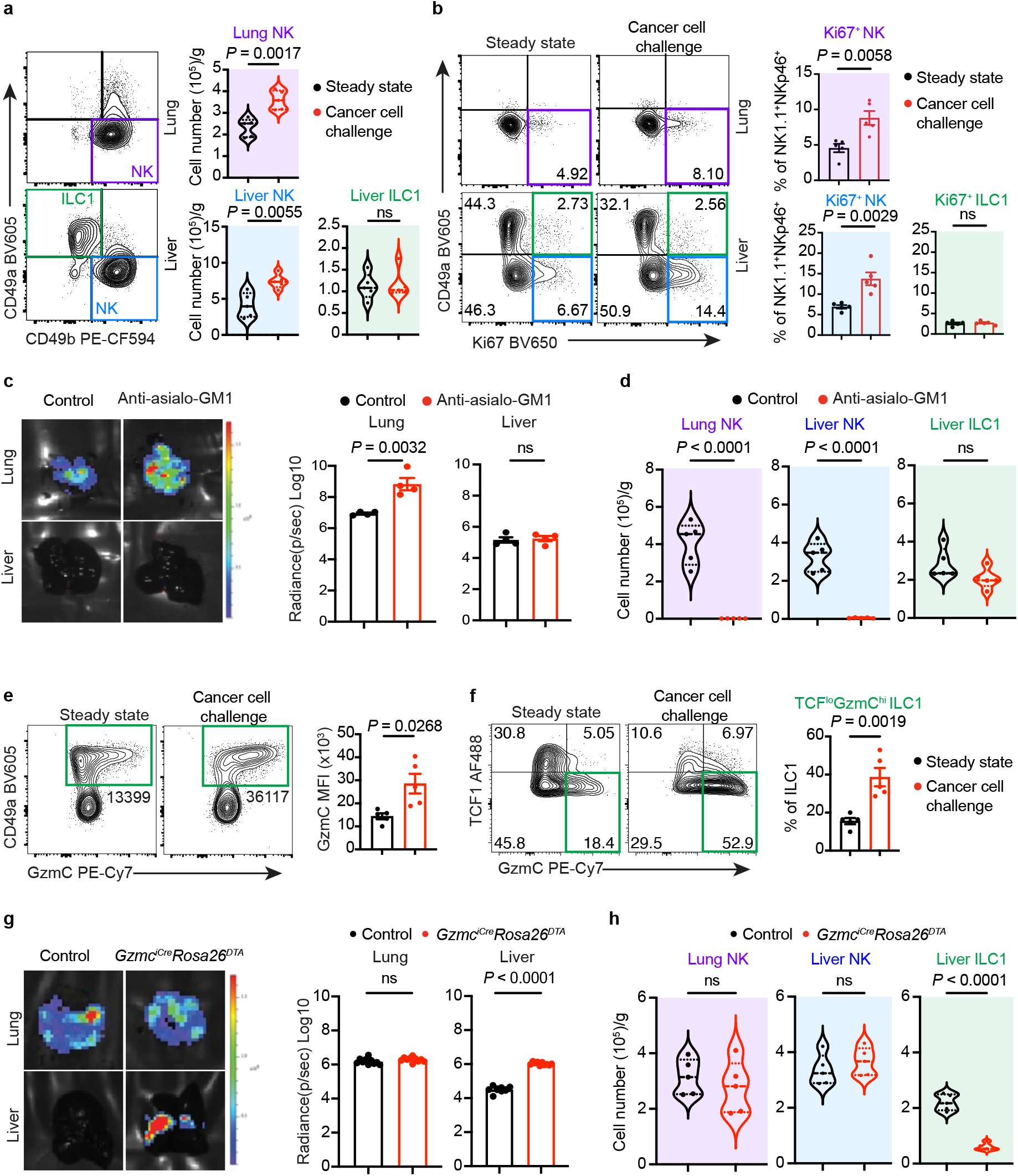
Hepatic ILC1s are selectively required for immunosurveillance of breast cancer liver metastasis. **a**, Representative flow cytometry analysis and quantification of NK1.1^+^NKp46^+^CD49a^+^CD49b^−^ ILC1s and NK1.1^+^NKp46^+^CD49a^−^CD49b^+^ conventional NK cells in the lungs and livers of C57BL/6 mice with or without E0771 cancer cell challenge for 24 h. Cell numbers per gram of tissue are shown as mean ± s.e.m. Mice were analyzed without tumor challenge. **b**, Representative flow cytometry analysis and quantification of Ki67 expression in ILC1s and NK cells in the lungs and livers of C57BL/6 mice with or without cancer cell challenge for 24 h. Percentages of Ki67^+^ cells among total group 1 innate lymphoid cells are shown as mean ± s.e.m. **c**, Representative bioluminescence images and radiance quantification of lung and liver colonization of cancer cells in C57BL/6 mice with or without anti-asialo-GM1 treatment (n = 5 per group). Data are representative of at least three independent experiments, each cohort is n = 5 per group, one representative image shown. Radiance quantification is shown as mean ± s.e.m. **d**, Quantification of ILC1 and NK cell numbers in the lungs and livers of C57BL/6 mice with or without anti-asialo-GM1 treatment. Lymphocyte numbers per gram of tissue are shown as mean ± s.e.m. Mice were analyzed without tumor challenge. **e**, Representative flow cytometry analysis and quantification of granzyme C (GzmC) expression in hepatic NK1.1^+^ NKp46^+^ group 1 innate lymphoid cells of C57BL/6 mice with or without cancer cell challenge. Quantification is shown as mean ± s.e.m. **f**, Representative flow cytometry analysis and quantification of the TCF1^lo^ GzmC^hi^ subset in hepatic ILC1s of C57BL/6 mice with or without cancer cell challenge. Quantification is shown as mean ± s.e.m. **g**, Representative bioluminescence images and radiance quantification of lung and liver colonization of cancer cells in *Gzmc^iCre^Rosa26^DTA^*and littermate control mice (n = 10 per group). One representative image shown. Data are representative of at least three independent experiments. Radiance quantification is shown as mean ± s.e.m. **h**, Quantification of ILC1 and NK cell numbers in the lungs and livers of *Gzmc^iCre^Rosa26^DTA^* and littermate control mice. Lymphocyte numbers per gram of tissue are shown as mean ± s.e.m. Mice were analyzed without tumor challenge. *P* values were calculated using Welch’s *t* test. ns: not significant (*P* > 0.05).

Despite failing to expand in response to cancer cell challenge, hepatic ILC1s expressed elevated levels of granzyme C (GzmC; Fig. 2e), a marker identifying activated ILC1s but not NK cells (Extended Data Fig. 6c)^19^. Within the ILC1 compartment, the more activated TCF1^lo^GzmC^hi^ subset, defined by GzmC upregulation and downregulation of the progenitor marker TCF1, was substantially enriched (Fig. 2f)^20^. To selectively deplete activated ILC1s, we generated *Gzmc^iCre^Rosa26^DTA^* mice^19^, which displayed significantly increased E0771.lmb colonization in the liver but not the lung (Fig. 2g), in association with markedly reduced hepatic ILC1s but an intact NK cell population (Fig. 2h, Extended Data Fig. 6d, e). Together, these findings reveal distinct, non-redundant functions of ILC1s and NK cells in restraining breast cancer liver and lung metastasis, respectively.

### DR5-mediated suppression of liver metastasis

Having established an essential role for ILC1s in the immunosurveillance of breast cancer liver metastasis, we next sought to define the underlying effector mechanisms. Under conditions of constitutive STAT5 activation, hepatic ILC1s can engage a perforin-dependent, lytic granule-mediated cytotoxic program that drives autoimmune pathology^19^. However, in mice challenged with E0771.lmb, perforin expression was upregulated in NK cells but not in ILC1s (Extended Data Fig. 7a). Consistent with this pattern, perforin deficiency resulted in increased lung, but not liver, metastasis (Extended Data Fig. 7b), indicating that the lytic granule pathway is required for NK cell-mediated control of pulmonary metastasis but dispensable for liver-specific surveillance mediated by ILC1s.

In addition to the lytic granule pathway, group 1 ILCs express the TNF superfamily member TRAIL, which has well-established metastasis-suppressive functions^21–24^. Genetic ablation of the TRAIL receptor DR5 (encoded by *Tnfrsf10b*) in mice likewise promotes cancer metastasis^25^. However, whether the metastasis-suppressive effects of TRAIL-DR5 signaling arise specifically from cancer cell-directed cytotoxicity remains unclear, as DR5 activation can trigger apoptosis in tumour-associated endothelial cells and immunosuppressive myeloid cells^26,27^, and cancer cell-autonomous DR5 signaling has also been reported to promote metastasis^28^. Notably, following E0771.lmb challenge, both NK cells and ILC1s upregulated TRAIL, with higher expression in ILC1s (Fig. 3a, Extended Data Fig. 7c).

**Fig. 3.**
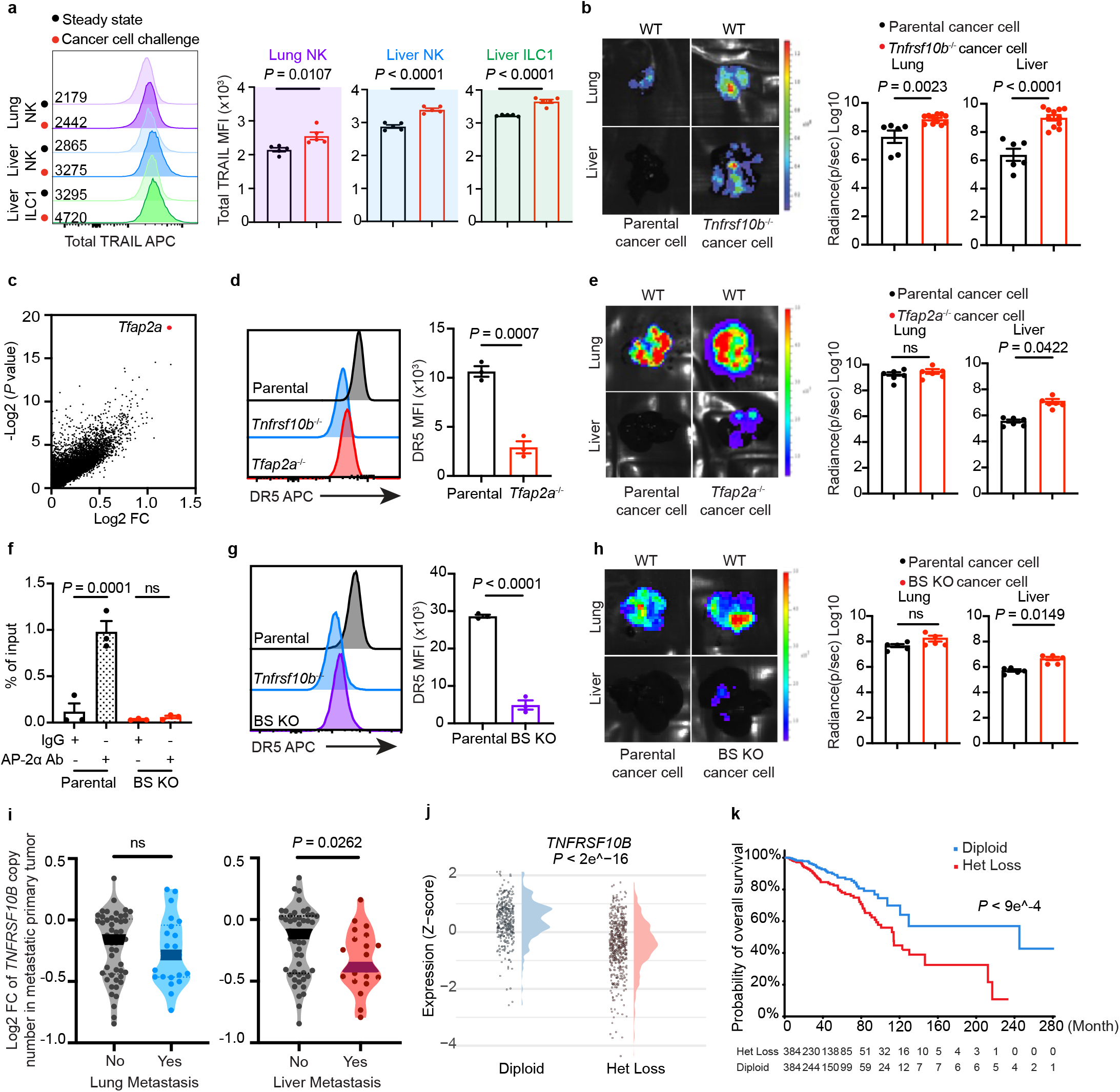
ILC1 surveillance of breast cancer metastasis is sensitive to DR5 expression, with *TNFRSF10B* heterozygous copy-number deletion predisposing patients to liver metastasis. **a,** Representative flow cytometry analysis and quantification of total (surface and intracellular) TRAIL expression in ILC1s and NK cells in the lungs and livers of C57BL/6 mice with or without E0771 cancer cell challenge for 24 h. TRAIL mean fluorescence intensity (MFI) is shown as mean ± s.e.m. **b**, Representative bioluminescence images and radiance quantification of lung and liver colonization of parental (n = 6) and DR5-deficient (*Tnfrsf10b*^−/−^; n = 11) E0771 cancer cells. One representative image shown. Data are representative of at least three independent experiments. Radiance quantification is shown as mean ± s.e.m. **c**, Identification of AP-2α (encoded by *Tfap2a*) as a critical regulator of DR5 expression in E0771 cancer cells using a CRISPR-Cas9 screen. Volcano plot showing Log_2_(fold change, FC) versus - Log_2_(*P* value) of genes identified by Illumina sequencing analysis. Each point represents an individual gene. Log₂ fold change (Log₂ FC) represents the enrichment or depletion of individual sgRNAs in the DR5-low cell population relative to their abundance in the unsorted library. Positive values denote sgRNAs enriched in DR5-low cells. **d**, Representative flow cytometry analysis and quantification of DR5 expression in parental and *Tfap2a*^−/−^ E0771 cancer cells. DR5 MFI is normalized to background signals from *Tnfrsf10b*^−/−^ E0771 cancer cells. **e**, Representative bioluminescence images and radiance quantification of lung and liver colonization of parental (n = 6) and AP-2α-deficient (*Tfap2a*^−/−^; n = 6) E0771 cancer cells. One representative image shown. Data are representative of at least three independent experiments. Radiance quantification is shown as mean ± s.e.m. **f**, Chromatin immunoprecipitation followed by qPCR (CHIP-qPCR) analysis of AP-2α binding to the *Tnfrsf10b* gene locus in parental E0771 cancer cells and in cells with knockout of the predicted AP-2α binding sites (BS KO) at the *Tnfrsf10b* gene locus. Enrichment of DNA immunoprecipitated with anti-AP-2α relative to control IgG was quantified over input DNA. **g**, Representative flow cytometry analysis and quantification of DR5 expression in parental and BS KO E0771 cancer cells. DR5 MFI is normalized to background signal from *Tnfrsf10b*^−/−^ E0771 cancer cells. **h**, Representative bioluminescence images and radiance quantification of lung and liver colonization of parental (n = 5) and BS KO (n = 5) E0771 cancer cells. One representative image shown. Data are representative of at least three independent experiments. Radiance quantification is shown as mean ± s.e.m. **i**, *TNFRSF10B* gene copy number in breast cancer patients with or without liver or lung metastases. Violin plots show log_2_-transformed fold changes in copy number, with each dot representing an individual patient sample and the horizontal line indicating the median. **j**, Association between *TNFRSF10B* gene copy number and mRNA expression. Expression levels of *TNFRSF10B* (z-score) are compared between samples with diploid or heterozygous deletion (het loss) of *TNFRSF10B*. **k**, Breast cancer patient survival stratified by *TNFRSF10B* copy number status (diploid versus het loss). *P* values were calculated using Welch’s *t*-test (**a**, **b**, **d**-**h**), Wilcoxon rank-sum test (**i**, **j**), and the log-rank (Mantel-Cox) test (**k**). ns: not significant (*P* > 0.05).

To evaluate cancer cell-intrinsic contributions of TRAIL-DR5 signaling to metastasis suppression, we generated DR5-deficient (*Tnfrsf10b^−/−^*) E0771.lmb cells using CRISPR-Cas9 gene editing (Extended Data Fig. 8a). Intravenous administration of *Tnfrsf10b^−/−^* E0771.lmb cells resulted in increased metastatic burden in both lung and liver compared with parental cells (Fig. 3b). This enhanced metastatic phenotype was preserved in *Rag1^−/−^*mice, which retain group 1 ILCs, but abolished in *Rag2^−/−^Il2rg^−/−^* mice lacking NK cells and ILC1s (Extended Data Fig. 8b, c). Depletion of DR5 from PY8119 *B2m^−/−^* cells also led to increased lung and liver metastasis (Extended Data Fig. 8d). Together, these findings argue against a cancer cell-autonomous role for DR5 in metastasis regulation and instead indicate that cancer cell DR5 is required for both NK cell-mediated control of lung metastasis and ILC1-mediated surveillance of liver metastasis across breast cancer cell lines.

### Partial DR5 loss as a liver metastasis risk factor

Having identified DR5 as a crucial breast cancer cell “alarmin” underlying NK cell- and ILC1-mediated metastasis immunosurveillance, we next sought to define the mechanisms governing DR5 gene regulation and their impact on lung and liver metastasis. To this end, we performed a genome-wide CRISPR-Cas9 screen in E0771.lmb cells based on perturbations of DR5 cell-surface expression (Extended Data Fig. 9a) and identified the transcription factor AP-2α (encoded by *Tfap2a*) as a positive regulator (Fig. 3c, Extended Data Fig. 9b, Extended Data Table 2). Using *Tnfrsf10b^−/−^* E0771.lmb cells as a knockout control, DR5 expression was reduced, but not abolished, in *Tfap2a^−/−^*cells (Fig. 3d). Notably, intravenous administration of *Tfap2a^−/−^*E0771.lmb cells resulted in substantially increased liver, but not lung, metastasis (Fig. 3e).

AP-2α, the founding member of the AP-2 transcription factor family, binds to the conserved 5’-GCCN_3_CCG-3’ motif at target gene loci and plays key roles in regulating cell proliferation and differentiation during vertebrate embryogenesis^29^, although its role in cancer remains ambiguous^30^. Analysis of the *Tnfrsf10b* promoter and enhancer regions revealed two candidate AP-2α binding sites (Extended Data Fig. 9c). To interrogate their function, we generated a binding-site knockout (BS KO) E0771.lmb line using CRISPR-Cas9 gene editing (Extended Data Fig. 9d, e). Chromatin immunoprecipitation followed by qPCR (ChIP-qPCR) confirmed AP-2α binding at the *Tnfrsf10b* gene locus in parental but not BS KO cells (Fig. 3f). Lack of these AP-2α binding sites diminished DR5 expression in BS KO cells, similar to that in *Tfap2a^−/−^*cells (Fig. 3d, g). Intravenous administration of BS KO cells increased liver, but not lung, metastasis (Fig. 3h), recapitulating the phenotype of *Tfap2a^−/−^* cells and contrasting with the broader metastasis enhancement observed in *Tnfrsf10b^−/−^* cells (Fig. 3b, e). To further characterize DR5 sensitivity for ILC1- and NK cell-mediated metastasis immunosurveillance, we isolated hepatic ILC1s and NK cells, and cultured them with parental, *Tnfrsf10b^−/−^*, or *Tfap2a^−/−^* E0771.lmb cells. While total DR5 deficiency similarly attenuated ILC1- and NK cell-induced cancer cell cytotoxicity, partial DR5 loss in *Tfap2a^−/−^* cells more substantially affected ILC1- than NK cell-mediated killing (Extended Data Fig. 9f). These findings establish *Tnfrsf10b* as a direct AP-2α target gene and reveal differential DR5 dose sensitivity for ILC1- and NK cell-mediated metastasis immunosurveillance.

To explore whether quantitative variation in DR5 expression influences metastasis organotropism in patients with breast cancer, we analyzed the MSK-IMPACT dataset. Although *TNFRSF10B* was not among the 505 target genes profiled for mutation and copy-number alterations in the standard MSK-IMPACT panel, partial loss of *NKX3-1*, the nearest profiled gene, located ∼600 kb from *TNFRSF10B* on chromosome 8, was associated with liver, but not lung, metastasis, whereas lung metastasis was associated with amplification of *MYC* and adjacent gene loci (Extended Data Fig. 10a). Subtype-stratified analysis revealed that copy-number loss at the *NKX3-1* locus was more strongly associated with liver metastasis in HR+ than TN or HER2+ breast cancer (Extended Data Fig. 10b). To directly assess whether partial *TNFRSF10B* copy-number loss predisposes to liver dissemination, we examined a subset of patients with breast cancer in MSK-IMPACT who underwent whole-exome sequencing. In this cohort, patients with liver, but not lung, metastasis were significantly more likely to harbor *TNFRSF10B* copy-number loss, transitioning from biallelic to monoallelic status (Fig. 3i). Analysis of breast tumours in TCGA further showed that heterozygous *TNFRSF10B* deletions reduced gene expression relative to diploid counterparts (Fig. 3j). Heterozygous *TNFRSF10B* loss also correlated with worse overall survival (Fig. 3k), consistent with the poor prognosis of patients with liver metastasis (Fig. 1b). Together, these findings support a tumour-suppressor function of DR5 and identify *TNFRSF10B* heterozygous copy-number loss as a risk factor for breast cancer liver metastasis.

### Rescuable ILC1 defects in steatotic livers

Following the observation that increased liver metastasis in HFD-fed mice was associated with selective depletion of group 1 ILCs (Fig. 1h), we further quantified hepatic NK cells and ILC1s, using lung tissue as a control. As expected, lung NK cells were comparable between HFD-fed and CD-fed mice; however, hepatic ILC1s, but not NK cells, were substantially reduced in HFD-fed mice (Fig. 4a, Extended Data Fig. 11a). Moreover, GzmC upregulation and expansion of the activated TCF1^lo^GzmC^hi^ ILC1 subset were impaired in HFD-fed mice following cancer cell challenge (Fig. 4b, Extended Data Fig. 11b), a finding validated by enumerating hepatic GzmC- YFP-reporter-positive CD3-ILC1s in *Gzmc^iCre^Rosa26^YFP^* mice (Extended Data Fig. 11c). Consistent with these observations, expression of hepatic ILC1, but not NK cell, signature genes was lower in steatotic relative to non-steatotic regions in human liver samples (Extended Data Fig. 11d, e). Together, these findings suggest that compromised ILC1 responses underlie the heightened susceptibility of steatotic livers to breast cancer metastasis.

**Fig. 4.**
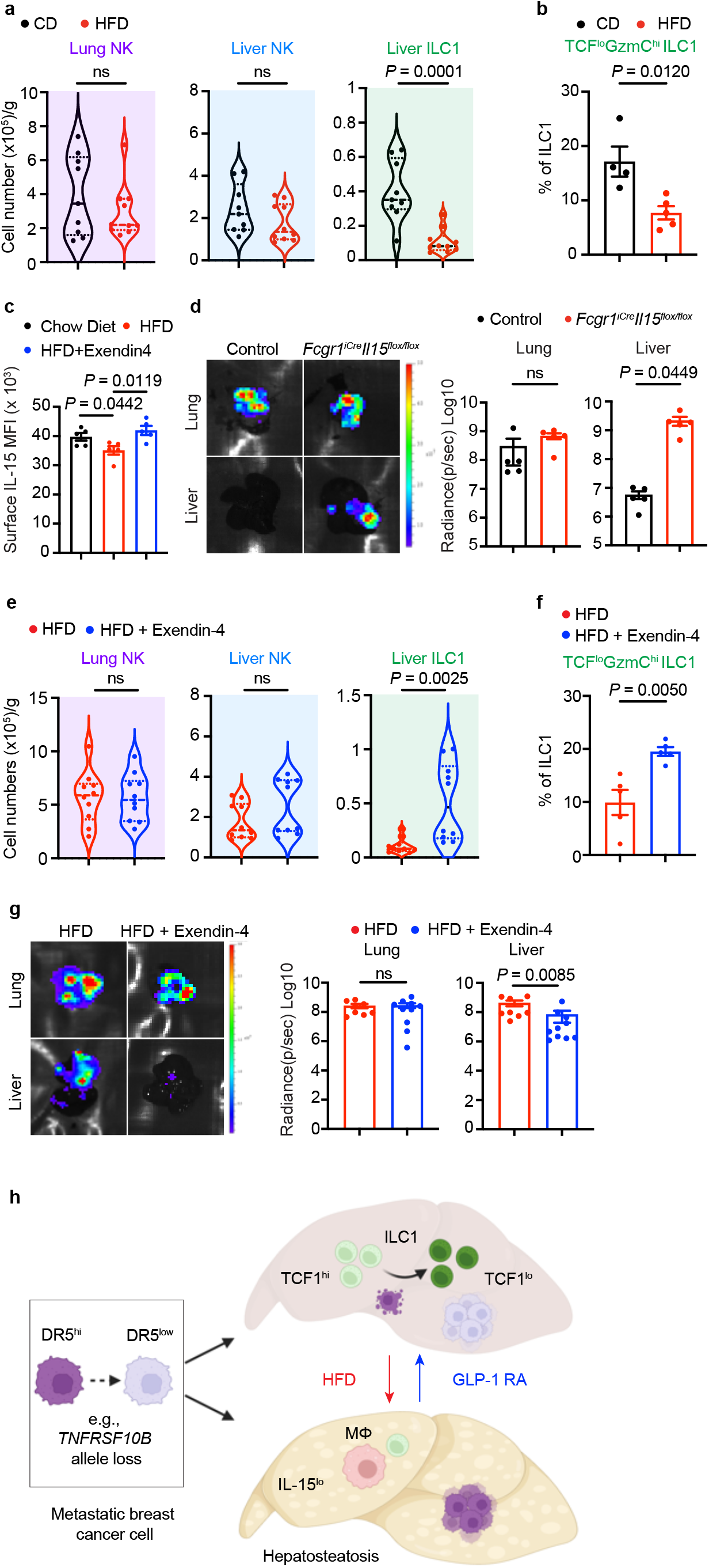
High-fat diet-induced ILC1 depletion and suppression, as well as breast cancer liver metastasis, are rescued by GLP1 receptor agonist treatment. **a**, **e**, Quantification of ILC1s and NK cell numbers in the lungs and livers of C57BL/6 mice fed a control chow diet (CD; n = 9) or high-fat diet (HFD; n = 10) (**a**), or HFD with (n =10) or without (n = 10) Exendin-4 treatment (**e**). Lymphocyte numbers per gram of tissue are shown as mean ± s.e.m. Mice were analyzed without tumor challenge. **b**, **f**, Quantification of TCF1^lo^GzmC^hi^ ILC1s in the livers of C57BL/6 mice fed with a CD (n = 4) or HFD (n = 5) (**b**), or HFD with (n = 5) or without (n = 5) Exendin-4 treatment (**f**). All mice were analyzed before tumour challenge. Percentages of TCF1^lo^GzmC^hi^ cells among ILC1s are shown as mean ± s.e.m. Mice were analyzed without tumor challenge. **c**, Quantification of cell-surface IL-15 expression in hepatic macrophages (MHCII^+^TIM4^+^) from CD (n = 5), HFD (n = 5), or HFD with Exendin-4 treatment (n = 5) mice. IL-15 MFI is shown as mean ± s.e.m. Mice were analyzed without tumor challenge. **d**, Representative bioluminescence images and radiance quantification of lung and liver colonization of E0771 cancer cells in control (n = 5) and *Fcgr1^iCre^Il15^flox/flox^* (n = 5) mice. One representative image shown. Radiance quantification is shown as mean ± s.e.m. **g**, Representative bioluminescence images and radiance quantification of lung and liver colonization of E0771 cancer cells in C57BL/6 mice fed an HFD with (n = 10) or without (n = 5) Exendin-4 treatment following cancer cell challenge. One representative image shown. Radiance quantification is shown as mean ± s.e.m. **h**, Schematic model illustrating how metastatic breast cancer cells evade hepatic ILC1-mediated immunosurveillance through cancer cell–intrinsic or microenvironmental mechanisms. In healthy liver, abundant ILC1s are activated in response to disseminated cancer cells and provide local innate immune protection. Metastatic breast cancer cells with high DR5 expression are effectively eliminated by ILC1s, whereas cells with reduced DR5 expression (e.g., through *TNFRSF10B* heterozygous allele loss) evade ILC1-mediated surveillance and survive in the liver. HFD-induced hepatosteatosis reduces IL-15 expression in hepatic macrophages, decreases hepatic ILC1 numbers, and compromises activated TCF1^lo^ ILC1 populations, driving metastatic outgrowth. GLP-1 receptor agonist (GLP-1 RA) treatment counteracts HFD-induced impairment of liver immunity, restoring ILC1-mediated surveillance and limiting liver metastasis. *P* values were calculated using Welch’s *t*-test. ns: not significant (*P* > 0.05).

Differentiation and function of group 1 ILCs, including tissue-resident ILC1s in cancer, are supported by the common gamma chain cytokine IL-15^31–33^. Hepatic ILC1s are maintained by IL-15 produced by parenchymal and stromal cell populations, including endothelial and myeloid cell lineages^34^. Notably, IL-15 expression was substantially reduced in liver macrophages, neutrophils, and eosinophils, but not endothelial cells, in HFD-fed mice relative to CD-fed mice, with macrophages expressing the highest levels (Fig. 4c, Extended Data Fig. 12a, b). To determine the function of macrophage IL-15, we crossed *Il15^flox^* mice onto the *Fcgr1^iCre^* background^35^ to selectively deplete IL-15 in macrophages, which led to reduced numbers of hepatic ILC1s but not NK cells (Extended Data Fig. 12c). This defect enhanced E0771.lmb cell metastasis to the liver but not the lung (Fig. 4d). Previous studies have shown that cancer cell metabolic reprogramming or myeloid cell accumulation could promote metastasis in steatotic livers^36,37^. To determine whether restoration of IL-15-dependent ILC1-mediated immunosurveillance would be sufficient to prevent breast cancer metastatic outgrowth, we fed a strain of IL-15 transgenic (*Il15^Tg^*) mice^38^ an HFD. IL-15 overexpression did not alter HFD-induced body weight gain or hepatic steatosis (Extended Data Fig. 12d, e). Strikingly, HFD-fed *Il15^Tg^* mice were protected from E0771.lmb cell metastasis to both lung and liver, accompanied by increased numbers of NK cells and ILC1s (Extended Data Fig. 12f, g). These findings suggest that defective ILC1 responses in HFD-fed mice are driven in part by a compromised macrophage IL-15 niche, with augmented IL-15 expression sufficient to expand ILC1s and suppress HFD-induced liver metastasis.

Diet-induced metabolic disorders such as type 2 diabetes and obesity can be effectively treated by glucagon-like peptide-1 (GLP-1) receptor agonists, which reprogram host metabolism^39^. Recent studies have also revealed that GLP-1 receptor agonists can revive NK cell effector functions independently of weight loss in patients with obesity^40^. To determine whether HFD-triggered hepatic ILC1 defects and steatotic metastasis could be corrected by these agents, we administered the GLP-1 receptor agonist Exendin-4 after 8 weeks of HFD feeding. Exendin-4 treatment significantly reduced body and liver weights (Extended Data Fig. 13a-c), and rescued IL-15 expression in macrophages (Fig. 4c, Extended Data Fig. 12b). Hepatic steatosis was also reduced, and hepatic ILC1 abundance, GzmC expression, and generation of the activated TCF1^lo^GzmC^hi^ ILC1 subset were restored, in Exendin-4-treated mice challenged with E0771.lmb cells (Fig. 4e, f; Extended Data Fig. 13d-f). Notably, Exendin-4 treatment effectively suppressed liver, but not lung, metastasis in HFD-fed mice across both the intravenous colonization model and the spontaneous fat-pad resection model (Fig. 4g, Extended Data Fig. 13g). These findings indicate that GLP-1 receptor agonist therapy not only reverses hepatic steatosis but also restores hepatic ILC1 functional competence, thereby enhancing immune-mediated suppression of liver metastasis.

## Discussion

Dissemination of cancer cells to distant organs is a heterogeneous and dynamically evolving process, with the liver representing not only the organ with the highest overall incidence of metastasis but also one associated with particularly fatal disease^7^. However, the basis of liver-specific metastatic susceptibility remains poorly defined. Herein, we identify hepatic ILC1s as a principal barrier constraining colonization of the liver by systemically disseminated breast cancer cells, in contrast to circulating NK cells, which primarily restrained lung colonization. These findings are consistent with prior evidence implicating group 1 ILCs in the control of breast cancer dormancy in the liver^41^, in which the respective contributions of ILC1s and NK cells were not distinguished. Moreover, recent studies demonstrating cooperative roles for ILC1s and NK cells in restraining liver metastasis of colon cancer cells disseminating through the portal vein^42–45^ suggest that cancer cell origin, metastatic burden, and/or route of hematogenous dissemination determine whether group 1 ILCs act selectively or collectively to suppress liver metastasis.

Our data also distinguish the characteristics and evasion mechanisms of ILC1 surveillance of breast cancer liver metastasis from those of NK cell-mediated suppression of lung metastasis. NK cells exhibit more potent immediate cytotoxicity than ILC1s^46^, and we find that they draw on both perforin- and DR5-dependent pathways to restrict lung colonization. Lung metastasis nonetheless remained incompletely controlled, which may reflect the migratory behaviour of mature NK cells and the brief window of their retention, recognition, and elimination of cancer cells within the lung microvasculature^47,48^. Liver-resident ILC1s, by contrast, have lower immediate cytotoxic potential but express higher levels of TRAIL and depend on high DR5 expression for effective cancer cell clearance. Consistent with this difference, monoallelic loss of *TNFRSF10B* (encoding DR5) was associated with liver, but not lung, metastasis in patients with breast cancer. Whether other oncogenomic or epigenetic alterations, including those affecting the AP-2α-DR5 axis described here, reduce DR5 expression sufficiently to evade ILC1-mediated hepatic immunosurveillance remains to be determined.

We further demonstrate that host-dependent evasion can arise from metabolic dysfunction: hepatic steatosis depleted hepatic ILC1s and impaired their activation, thereby weakening a critical organ-specific barrier to disseminated breast cancer cells. Notably, BMI (hepatic steatosis)- and *TNFRSF10B* copy-number deletion-associated liver metastasis risk was strongest in TN and HR+ patients, respectively, suggesting that the relative weight of host metabolic states and cancer genomic features as liver metastasis susceptibility factors differs by breast cancer subtype. Importantly, restoration of ILC1 abundance and function through IL-15 augmentation or GLP-1 receptor agonism underscores that the immune-disruptive hepatic metabolic state is a tunable determinant of metastatic susceptibility. Together, these findings establish ILC1s as central orchestrators of liver-specific immunosurveillance and illustrate how cancer cell–intrinsic alterations and host metabolic perturbations converge to shape organotropic metastasis. Our study also identifies actionable therapeutic entry points, including GLP-1 receptor agonists, to rectify hepatic steatosis-associated ILC1 defects and restore resistance to breast cancer liver metastasis (Fig. 4h). More broadly, this work advances the concept that tumour–immune crosstalk is profoundly shaped by organ-specific metabolic and immunological contexts, opening new opportunities to intercept metastasis at its earliest stages.

## Methods

### Patient cohort and data analysis

Breast cancer cases with multi-organ metastasis were drawn from an MSK-IMPACT Clinical Sequencing Cohort. Patient-level genomic and clinical data were obtained from the MSK investigator-accessible cBioPortal resource (https://cbioportal.mskcc.org) under the MSK-IMPACT study (Study ID: mskimpact; accessed August 19, 2025). All analyses followed MSK data-use policies and MH RDG requirements. For each cancer patient with multiple metastatic organ sites, the latency was defined as the time from the first-ever metastasis to the organ-specific metastasis diagnosis. Metastatic events were classified as Early (≤ 365 days) or Late (> 365 days) based on this interval. Patients exhibiting both early and late events across different sites were selected for further analysis. Each combination of organ site and timing (e.g., Liver_Early, Bone_Late) was treated as a distinct pattern, and a binary incidence matrix was constructed using the UpSetR package in R. Patient steatosis was assessed by review of available liver histology and radiologic studies. For patients with liver biopsy specimens containing evaluable background liver parenchyma, steatosis was graded according to the NIH NASH Clinical Research Network (CRN) criteria^49^. For all patients, available radiology reports, including CT, MRI, and ultrasound, were reviewed for documentation of hepatic steatosis. Patients without available or documented imaging information were classified as undetermined rather than as having no steatosis.

For metastatic site analysis of *TNFRSF10B* (encoding DR5) copy number, we analyzed breast cancer cases from the MSK-IMPACT clinical recaptured WES cohort (whole-exome recapture and resequencing of IMPACT tumours). Patient-level genomic and clinical data were obtained from the MSK investigators accessible cBioPortal resource (https://cbioportal.mskcc.org) under the MSK-IMPACT Clinical Recaptured WES Cohort (Study ID: mixed_impact_wes_2017; accessed on September 20, 2025). All analysis followed MSK data-use policies and MH RDG requirements. Gene copy number values were extracted from segmented copy number data. For each metastatic site, samples were grouped by the presence or absence of organ-specific metastasis, and differences in copy number were assessed using the Wilcoxon rank-sum test. For *TNFRSF10B* copy number versus expression level and survival analyses, patient data were obtained from the TCGA breast cancer carcinoma dataset and analyzed in MSK cBioPortal.

Human liver Visium datasets (MASLD and healthy controls) were reported previously (GEO: GSE192741)^50^. Data from 5 samples from 4 patients with varying degrees of hepatic steatosis were downloaded as filtered feature-barcode matrices (i.e., the output of Space Ranger as uploaded by Guilliams *et al.*) together with corresponding histological images, and processed in R using Seurat (v5.2.1). Following merging of samples into a single Seurat object, dimensionality reduction of Visium spots via principal component analysis (PCA) was performed using 2,000 highly variable genes. Spots were subsequently clustered using the Louvain algorithm (resolution 1.1) on a 30-nearest-neighbor graph. One cluster was noted via differential expression analysis to be highly enriched for genes involved in lipid metabolism and, upon comparison with hematoxylin and eosin–stained sections from each sample, found to correspond to areas annotated as steatotic. Expression of manually curated hepatic ILC1 and NK gene signatures was calculated for each Visium spot using the AddModuleScore function in Seurat. The ILC1 signature included the following genes: *ZNF683*, *ITGA1*, *CXCR6*, *CD69*, *CD200R1*, *NCR3*, *KLRD1*, *CD52*, *KLRC1*, *KLRC2*, and *TNFSF10*. The NK cell signature included the following genes: *B3GAT1*, *CX3CR1*, *FCGR3A*, *KIR2DL3*, *KIR3DL1*, *KLRF1*, *LAIR2*, *MTSS1*, *S1PR5*, and *SELL*^51^. Signature expression was statistically compared between steatotic regions (i.e. spots assigned to the steatotic cluster) and non-steatotic regions (all spots assigned to other clusters) using two-sided t-tests and visualized as box plots.

### Mice

C57BL/6, *Rag1^−/−^*, *Rag2^−/−^Il2rg^−/−^*, *Prf1^−/−^*, *Rosa26^DTA^*, and *Rosa26^YFP^* mice were purchased from The Jackson Laboratory. *Il15*^flox^, *Il15^Tg^*, *Gzmc^iCre^*, *Fcgr1*^iCre^, and *Ncr1^iCre^* mice were previously described^15,17,19,35,38^, and were generously provided by colleagues or made in the M.O.L. lab. For high-fat diet or calorie-matched control diet treatment, mice were fed a 60% kcal Fat (Research Diets, Cat. #D12492i) or a 10% kcal Fat (Research Diets, Cat. #D12450J) for 16 weeks before experimentation. All mice used in these studies were on the C57BL/6 background and maintained under specific pathogen-free conditions. Animal experimentation was conducted in accordance with procedures approved by the Institutional Animal Care and Use Committee of Memorial Sloan Kettering Cancer Center (MSKCC). Mice aged 6–10 weeks were used for experiments unless otherwise noted.

### Cell lines

The E0771.lmb breast cancer cell line (ATCC, CRL-3405), PY8119 breast cancer cell line (CRL-3278) stably transduced with a luciferase-expressing vector pHIV-Luc-ZsGreen (Plasmid #39196, Addgene) was generated to facilitate the tracking of metastasis. *Tnfrsf10b^−/−^*, *Tfap2a ^−/−^*, AP-2α binding-site knockout (BS KO) cells, and *B2m^−/−^* were generated using CRISPR-Cas9 gene editing. Briefly, E0771.lmb or PY8119 cells were collected and extensively washed with PBS. 4 × 10^5^ cells were electroporated with recombinant Cas9 protein v.3 (IDT) complexed with gene locus–specific sgRNA using Lonza 4D-nucleofector unit and buffer kit following manufacturer’s instructions. 2.5 μg sgRNA and 5 μg Cas9 recombinant protein were used per 100 μl reaction. The following sgRNA sequences were used: *Tnfrsf10b*, 5’- GCATCGACACACCGTATTTG-3’ and 5’- TAGGCCAGCTGGACGATTAT-3’; *Tfap2a*, 5’-AGCGACGCGTTAAGACACTC-3’ and 5’- GGACAACCTCTTCGGTGGCG-3’; BS target, 5’-GCTGACTCAAACGTCCCTAC-3’ and 5’- GATGTGGTAGCATGCGACAT-3’; *B2m*, 5’-CACGCCACCCACCGGAGAAT-3’ and 5’- CTGGTGCTTGTCTCACTGAC-3’. Following electroporation, *Tnfrsf10b*^−/−^ and *B2m^−/−^* cells were sorted by flow cytometry based on DR5 and H-2K^b^ surface staining, respectively. *Tfap2a*^−/−^ and BS KO cells were seeded in 96-well plates for single-cell clone selection. Target gene deletion was validated by immunoblotting and genotyping. All cells were cultured in Dulbecco’s Modified Eagle Medium (DMEM) high-glucose medium (Media Preparation Core, MSKCC, Cat. #52100047). Medium was supplemented with 10% fetal bovine serum (FBS) (Cytiva HyClone, Cat. #SH30910.03), 2 mM L-glutamine (Thermo Fisher Scientific, Cat. #25030081), 100 IU/mL penicillin-streptomycin (Thermo Fisher Scientific, Cat. #15140163). All cells were grown in a humidified incubator at 37°C with 5% CO_2_ and verified to be mycoplasma-free.

### Metastasis models

#### Mammary fat pad implantation, primary tumour resection, and longitudinal metastasis monitoring

E0771.lmb cells were harvested and washed with PBS. Mice were anaesthetized with 2-3% isoflurane in oxygen, placed in dorsal recumbency on a warming pad and given pre-operative analgesia. The fourth mammary fad pad (inguinal) was exposed through a 4 mm skin incision. Using a 29-gauge insulin syringe, 2.5 × 10^5^ cells in 50 μl PBS were injected into the cleared fat pad. The skin was closed with absorbable suture. When tumours reached 200-250 mm^3^ (12-15 days post-implantation), mice were re-anaesthetized with isoflurane. The previous incision site was reopened, and the tumour was bluntly dissected free of surrounding tissue. Metastatic dissemination was tracked by *in vivo* bioluminescence imaging using an IVIS Spectrum system (Caliper Life Sciences).

#### Intravenous inoculation of cancer cells to model systemic metastasis dissemination

5 × 10^5^ E0771.lmb, PY8119 *B2m*^−/−^ cancer cells or their derivatives were collected and washed with PBS. Cells were resuspended in 100 µL ice-cold DPBS (Dulbecco’s phosphate-buffered saline) without calcium or magnesium (Thermo Fisher Scientific, Cat. #14190250) and transferred to mice via tail vein injection. Metastasis burden was monitored and quantified by *in vivo* bioluminescence imaging using an IVIS Spectrum system (Caliper Life Sciences).

For bioluminescence imaging, mice were injected with freshly prepared D-luciferin (150 mg/kg) via the retro-orbital route 5 min before euthanasia. Lungs and livers were dissected and imaged individually. Luciferase signals were quantified using Living Image software (Caliper Life Sciences).

### Antibody treatment

Anti-NK1.1 antibody was isolated from PK136 hybridoma cells (ATCC, HB-191) and administered via intravenous injection at 200 µg/mouse three days before inoculation of E0771.lmb cells. Anti-asialo-GM1 antibody (Wako Chemical USA, Cat. #NC1345696) was administered intraperitoneally twice weekly at 10 µg/mouse.

### Immunofluorescence and Oil Red O staining

Liver tissues were collected from mice, washed twice with PBS, fixed in 4% periodate-lysine-paraformaldehyde for 16–24 h, dehydrated in 30% sucrose for 24 h, and then snap-frozen in OCT. Sections were cut at 20 µm thickness, blocked for 60 min in staining buffer (0.1 M Tris, 1% BSA, 1% FBS, 0.3% Triton X-100), and stained with fluorescently conjugated antibodies overnight at 4°C. Slides were washed and counterstained with DAPI (4’,6-diamidino-2-phenylindole; Sigma), mounted with Fluoromount-G (Invitrogen, Cat. #00-4958-02) and dried overnight in at 4°C. Images were captured using a Mirax Scan system (Zeiss) with four-color channels. For Oil Red O staining, the section slides were stain using Mercedes Scientific Oil Red O stain kit (Mer ORK2). Images were captured using a Mirax Scan system (Zeiss) with bright field and quantified using Image J.

### Hematoxylin and eosin staining

Liver tissues were collected from mice, washed twice with PBS, fixed in 4% periodate-lysine-paraformaldehyde for 16–24 h. After fixation, samples were rinsed three times in ice-cold PBS. Fixed tissues were dehydrated through a graded ethanol series (70%, 80%, 95%, and 100%; 30 min each) followed by two changes of 100% ethanol (45 min each). Samples were cleared in xylene twice for 20 min, infiltrated with 1:1 xylene–paraffin (60°C, 30 min) and then with molten paraffin (Paraplast Plus; Sigma-Aldrich) in a vacuum oven at 60°C for three changes (45 min each). Tissues were oriented in fresh paraffin and allowed to solidify at room temperature. Paraffin blocks were stored at 4°C until sectioning. Blocks were trimmed and serially sectioned at 5 µm thickness. Ribbons were floated on a 42°C water bath containing 0.01% Tween-20, collected onto Superfrost Plus slides (Thermo Fisher Scientific) and dried overnight at 37°C. Prior to staining, slides were baked at 60°C for 1 h to enhance adhesion and remove residual paraffin. Slides were deparaffinized in two changes of xylene (10 min each), transferred through a descending ethanol series (100%, 95%, 80%, 70%; 2 min each) and rinsed in running distilled water for 5 min. Sections were stained in Mayer’s hematoxylin (Sigma-Aldrich) for 6 min, differentiated in 0.3% acid alcohol (1% HCl in 70% ethanol, 5 s) and “blued” in running tap water for 5 min. Slides were counterstained in 1% eosin Y (aq.) for 45 s, rapidly dehydrated through 95% and 100% ethanol (2 × 30 s each) and cleared in xylene (2 × 2 min). Images were captured using a Mirax Scan system (Zeiss) with bright field.

### Cell isolation from tissues

Euthanized mice were perfused with 10 mL of 2 mM EDTA in PBS via the portal vein or right ventricle. Cells were isolated from the liver and lung as follows. Tissues were manually disrupted with a razor blade, digested with 1 mg/mL collagenase D (Sigma-Aldrich, Cat. #C5138) and 4 µg/mL DNase I (Sigma-Aldrich) in RPMI-1640 at 37°C for 30 min with vortexing every 10 min. Digested tissues were passed through a 70 μm strainer. To enrich for lymphocytes, liver and lung cell pellets were resuspended in 44% Percoll and layered above 66% Percoll, centrifuged at 1,900 × g for 30 min at room temperature without brake. Cells at the interface were collected, stained, and analyzed by flow cytometry. For isolation of myeloid cells from the liver, a previously published protocol was followed^52^.

### Flow cytometry

For flow cytometry, cells were first stained with Zombie NIR^TM^ Fixable Viability Kit (BioLegend) in PBS for 15 min at 4°C to identify dead cells. For cell-surface staining, cells were incubated with an antibody cocktail on ice in the presence of anti-CD16/CD32 antibodies in BD Horizon^TM^ Brilliant Staining Buffer for 30 min. For intracellular staining, the Tonbo or Thermo Fisher Scientific Transcription Factor Fix/Perm Kit was used according to the manufacturer’s instructions. All samples were acquired on an Aurora cytometer (Cytek) using the SpectroFlo software (v2.2.0.3) and analyzed with FlowJo (v10.6.1).

### CRISPR/Cas9 screen

The parental E0771.lmb cells were transduced with a lentivirus vector driving expression of the Cas9 protein. The resulting E0771-Cas9 cell line was transduced with a Mouse CRISPR Knockout Pooled Library (GeCKO v2 with combined #1000000052 and #1000000053 pooled lentiviral libraries from Addgene). A total 2 × 10^8^ cells were infected with the sgRNA lentivirus library. Twelve hours after infection, medium was replaced with fresh medium containing 5 μg/ml puromycin to eliminate non-infected cells. After 7 days, 5 × 10^8^ cells were stained with biotinylated DR5 antibody (R&D Systems, Cat. #BAF721) for 30 min on ice, washed once with PBS, and then stained with streptavidin-APC (Thermo Fisher Scientific, Cat. #17-4317-82) for 30 min. Cells were washed twice with PBS, and the top and bottom 15% of cells by DR5 signal intensity were sorted by flow cytometry; 1 × 10^8^ cells collected before DR5 staining served as a control. Genomic DNA was isolated using a DNA isolation kit (Zymo Research, Cat. #D4075). PCR of sgRNAs for Illumina sequencing was performed following the Broad Institute protocol^53^. For the DR5-low population, sgRNA enrichment fold change and FDR q-values were calculated using MAGECK^54^ with default settings.

### *In vitro* cytotoxicity assay

ILC1s and NK cells were isolated from the liver based on the protocol describe above. NK cells (CD45^+^TCRβ^−^NKp46^+^NK1.1^+^CD49b^+^CD49a^−^) and ILC1s (CD45^+^TCRβ^−^NKp46^+^NK1.1^+^CD49b^−^ CD49a^+^) were sorted by flow cytometry and co-cultured with luciferase-expressing target cancer cells at an effector-to target (E:T) ratio of 5:1 (5,000 effector cells and 1,000 target cells per well in 96-well plates) in the presence of 10 ng/ml IL-15/IL-15Rα complex. After 12 h, viable target cells were quantified by measuring luminescence following addition of D-luciferin (150 μg/ml). Cytotoxicity was calculated as the percentage reduction in luminescence compared to target cells cultured alone.

### Chromatin immunoprecipitation followed by qPCR (ChIP-qPCR)

Chromatin isolation and immunoprecipitation were performed as previously described^55^. Briefly, 1.5 × 10^7^ parental or BS KO E0771.lmb cells were harvested for immunoprecipitation with anti-AP-2( antibody (Invitrogen, Cat. #MAI-872) and rabbit IgG isotype control (Cell Signaling Technology, Cat. #3900S). ChIP-qPCR was performed using primers spanning the AP-2(-binding sites (5’-CAAATTCCTGTTCTTCATCCTGCGAG-3’; 5’-GTGGTGCGGGAAGGCATTGG-3’) and PowerUp SYBR Master Mix (Thermo Fisher Scientific, Cat. #A25777). Data were normalized to input and presented as percent input.

### Fatty acid and lipid profiling

Mouse plasma was extracted using a Folch extraction. To 20 µL of plasma, 30 µL of water was added, followed by the addition of 300 µL of methanol containing internal standards: 0.5 µM d-31 palmitate (Cayman Chemical) and 1 µg/mL EquiSPLASH (Avanti Research). After brief vortexing, 600 µL of chloroform was added. Thereafter, 180 µL of water was added to induce phase separation and incubated at room temperature for 10 min. After vortexing, the extract was centrifuged at 3000 g for 5 min at 4°C. Afterwards, the lower, chloroform layer was obtained and transferred to a glass vial. Samples were then re-extracted with 700 µL of chloroform:methanol (2:1 v/v). After brief vortexing, the lower, chloroform layer was collected and pooled with the previous chloroform layer in the same glass vial. The combined chloroform layer was split into equal volumes (500 µL) for fatty acid and lipid analysis respectively. The organic extracts were then dried under nitrogen and each reconstituted in 100 µL of acetonitrile:isopropanol (1:1 v/v) for LC–MS analysis.

Fatty acid profiling was performed using an Agilent 6546 Q-TOF mass spectrometer operated in negative ion mode, coupled to an ACQUITY UPLC BEH C18 column (100 mm × 2.1 mm, 1.7 µm; Waters). Mobile phase A consisted of 10 mM ammonium acetate in 60:40 acetonitrile:water, and mobile phase B consisted of 10 mM ammonium acetate in 85:10:5 isopropanol:acetonitrile:water. The LC gradient was as follows: 0 min, 10% B; 2 min, 10% B; 16 min, 60% B; 21 min, 100% B; 24.1 min, 10% B; 27.6 min, 10% B. Other LC parameters included a flow rate of 0.3 mL/min, a column temperature of 45°C, and an injection volume of 5 µL. MS source parameters included: gas temperature: 200°C; gas flow: 11 L/min; nebulizer pressure: 50 psig; sheath gas temperature: 300°C; sheath gas flow: 12 L/min; VCap: 3000 V; nozzle voltage: 0 V; fragmentor: 150V. Data were acquired from m/z 50–1700 with active reference mass correction (m/z 301.9981 and 980.0163).

Lipid profiling was performed using an Agilent 6546 Q-TOF mass spectrometer in both positive and negative ionization modes, coupled to a ZORBAX Eclipse Plus C18 column (100 mm × 2.1 mm, 1.8 µm particle size; Agilent Technologies). Mobile Phase A consisted of 10 mM ammonium formate in 50:30:20 water:acetonitrile:isopropanol. Mobile Phase B consisted of 10 mM ammonium formate in 1:9:90 water:acetonitrile:isopropanol. The LC gradient was as follows: 0 min, 0% B; 2.7 min, 45% B; 2.8 min, 53% B; 9 min, 65% B; 9.1 min, 89% B; 11 min, 92% B; 11.1 min, 100% B; 12 min, 10% B; 15 min, 10% B. The flow rate was 0.4 mL/min; column temperature was maintained at 60°C, and the injection volume was 5 µL. MS source parameters were as follows: gas temperature, 200°C; gas flow, 10 L/min; nebulizer pressure, 50 psig; sheath gas temperature, 300°C; sheath gas flow, 12 L/min; VCap, 3,000 V; nozzle voltage, 0 V; fragmentor, 150 V. Data were acquired from m/z 50–1700 with active reference mass correction (positive mode: m/z 322.0481 and 922.0097; negative mode: m/z 301.9981 and 966.0007). Five iterative MS/MS acquisitions were performed on pooled lipid extracts in both polarities for lipid annotation. Lipid identification performed using Agilent Lipid Annotator (v1.0.1). Peak picking and integration was performed using Skyline (MacCoss Lab).

### Antibodies and chemical compounds

The following antibodies were used in flow cytometry: BV570 anti-CD45 (30-F11, BioLegend, 1:200), BV605 anti-CD49a (Ha31/8, BD Biosciences, 1:200), PE-CF594 anti-CD49b (DX5, BD Biosciences,1:200), PE-Cy7 anti-GzmC (SFC1D8, BioLegend, 1:200), AF488 anti-TCF1 (S33-966, BD Biosciences,1:100), biotin anti-TRAIL (N2B2, Invitrogen, 1:200), biotin anti-DR5/TNFRSF10B (BAF721, R&D Systems, 1:400), FITC anti-CD8*_a_* (53-6.7, Cytek, 1:200), BV750 anti-CD4 (GK1.5, BioLegend, 1:200), BV650 anti-CD8β (53-5.8, BD Biosciences,1:200), BV480 anti-TCRβ (H57-597, BD Biosciences,1:200), APC anti-CD1d tetramer (NIH Tetramer Core Facility, 1:200), PerCP-eFluror 710 anti-TCRγδ (eBioGL3, Invitrogen, 1:200), PE-Cy5.5 anti-Foxp3 (FJK-16s, Invitrogen, 1:200), BV510 anti-CD19 (1D3, BD Biosciences, 1;200), BV711 anti-NKp46 (29A1.4, BioLegend, 1:200), Mouse Nkp46 antibody (AF2225, R&D systems), APC anti-NKp46 ( 29A1.4, BioLegend, 1:200), APC-Cy7 anti-NK1.1 (PK136, Cytek, 1:200), BV421 anti-GATA3 (L50-823, BD Biosciences, 1:200), BV786 anti-RORγ (Q31-378, BD Biosciences, 1:200), PE anti-Perforin (S16009A, BioLegend, 1:200), Biotin anti-mouse IL-15 (PeproTech, 1:500), PE Streptavidin (eBioscience, 1:500), Alexa Fluor 647 anti-mouse CD31(MEC13.3, Biolgend, 1:200), BV480 anti-TCRγδ (GL3, BD Biosciences, 1:200), BV480 anti-CD19 (1D3, BD Biosciences,1:200), BV711 anti-Ly6G (1A8, BD Biosciences,1:200), BV605 anti-MHC-II (M5/114.15.2, BD Biosciences, 1:200), PerCP-Cy5.5 anti-Siglec F (E50-2440, BD Biosciences, 1:200), BV786 anti-TIM-4 (RMT4-54, BD Biosciences, 1:200), BV421 anti-F4/80 (T45-2342, BD Biosciences, 1:200), APC-Cy7 anti-Ly6C (AL 21, BD Biosciences, 1:200), APC anti-mouse H-2Kb (AF6-88.5, BioLegend, 1:200). The following antibodies were used for immunofluorescence: Alexa Fluor 594 anti-CD3 (17A2, BioLegend, 1:200), Alexa Fluor 647 anti-CD38 (90, BioLegend, 1:200), Anti-GFP (Abcam, Cat. #ab13970, 1:1000), Alexa Fluor 488 anti-chicken IgY (Jackson ImmunoResearch, Cat. #703-545-155, 1:500). Exendin-4 (MedChemExpress, Cat. #HY-13443) was administered by daily intraperioneal injection at 24 nmol/kg.

### Statistical analysis

Statistical analyses were performed using GraphPad Prism software. Statistical details for all experiments are provided in the figure legends. Bar graphs display mean ± s.e.m., with each dot representing one biological replicate. *P* values are indicated on graphs or in figure legends.

## Acknowledgments

We thank the patients and families who contributed to the MSK-IMPACT clinical sequencing program. We are grateful to members of the Li lab and our collaborators for helpful discussions. We thank the Memorial Sloan Kettering Cancer Center Flow Cytometry Core, Integrated Genomics Operation Core, Cell Metabolism Core and Molecular Cytology Core for technical assistance. This work was supported by a National Institutes of Health grant (R01CA243904), a Memorial Sloan Kettering Cancer Center Core Grant (P30 CA008748), and postdoctoral fellowships from the Center for Experimental Immuno-Oncology and the Ludwig Center for Cancer Immunotherapy at Memorial Sloan Kettering Cancer Center.

## Author Contributions

P.L. and M.O.L. were involved in all aspects of this study, including conception, design, and performance of experiments, data analysis and interpretation, and manuscript writing. J.L. performed patient data analysis from the MSK-IMPACT and TCGA databases. A.E.C. analyzed the human liver Visium dataset. D.B. and F.P. analyzed the patient liver histopathology and radiology reports. J.Z., X.Z., Z.X., L.J., S.G., S.B., and B.G.N. assisted with experiments and mouse colony management. J.Y. and P.C. assisted with ChIP-qPCR experiments. P. R., C.F., A.P., and A.R. provided access to the MSK-IMPACT dataset. All authors provided critical feedback on the results and manuscript.

## Extended data figure legends

**Extended Data Fig. 1.**
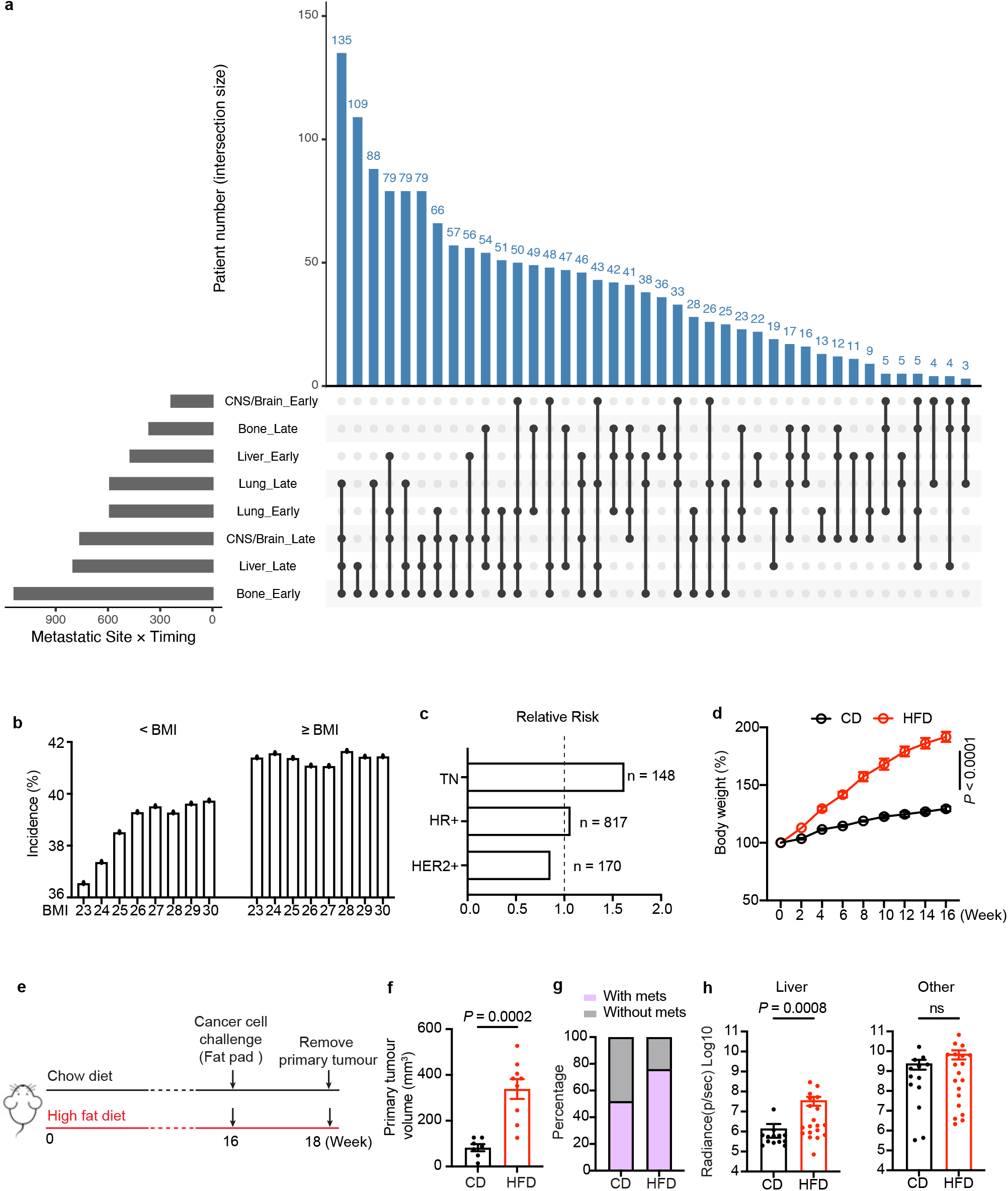
Temporal and organ-specific distribution of breast cancer metastases in patients and in an orthotopic mouse model. **a**, UpSet plot showing the number of patients exhibiting distinct combinations of organ-specific temporal patterns of metastasis. Each bar represents the number of patients sharing a particular intersection of early (≤ 1 year after first metastasis) and late (> 1 year after first metastasis) metastatic events across major organs (liver, lung, bone, and CNS/brain). Filled dots along the bottom matrix indicate which organ-timing categories are included in each intersection, and vertical bars above quantify the corresponding patient counts. **b**, Impact of varying Body Mass Index (BMI) stratification thresholds on the incidence of liver metastasis in breast cancer patients. Incidence rates of liver metastasis across a dynamic range of clinical BMI cut-off thresholds spanning from 23 to 30 kg/m². For each indicated threshold, the metastatic incidence was calculated independently for the low BMI cohort (below the cut-off value) and the high BMI cohort (equal to or above the cut-off value). The incidence percentage represents the absolute number of patients diagnosed with confirmed liver metastasis divided by the total number of patients within that specific stratified sub-population. **c**, Subtype-specific Relative Risk (RR) of liver metastasis associated with elevated BMI ( ≥ 23 kg/m²) in breast cancer patients. RR of developing liver metastasis in patients with elevated body mass index (BMI ≥ 23 kg/m²) compared to the baseline control cohort (BMI < 23 kg/m²), stratified by major clinical molecular subtypes: Triple-Negative (TN), Hormone Receptor-Positive (HR+), and HER2-Amplified (HER2+). The RR for each independent subtype was calculated as the ratio of the metastatic incidence percentage within the high-BMI group ( ≥ 23 kg/m²) to the metastatic incidence percentage within the low/normal-BMI reference group (< 23 kg/m²). An RR value greater than 1.0 indicates an increased risk of liver metastasis associated with a high BMI. **d**, Relative body weight change over time in diet groups. Biweekly body weights of mice fed a control chow diet (CD; n = 10) or high-fat diet (HFD; 60 kcal% fat; n = 10) at the indicated timepoints. Mice were started on their respective diets at 8 weeks of age. **e**, Schematic workflow illustrating the orthotopic mammary fat pad injection, primary tumour resection, and subsequent monitoring of metastasis progression. **f**, Quantitative analysis of primary tumour volume following orthotopic fat pad injection in mice maintained on either a CD (n = 7) or HFD (n = 9). **g**, Quantification analysis showing the proportion of mice with confirmed metastasis after surgical removal of the primary tumour in the CD or HFD groups (n = 25 per group). **h**, Quantification of metastatic lesions detected in the liver and other sites. Other sites evaluated include the lung, intestine, brain, and the abdominal region (including general abdominal tissue and abdominal white adipose tissue) *P* values were calculated using two-way ANOVA (**d**), Mann-Whitney *U* test (**f, h**), ns: not significant (*P* > 0.05).

**Extended Data Fig. 2.**
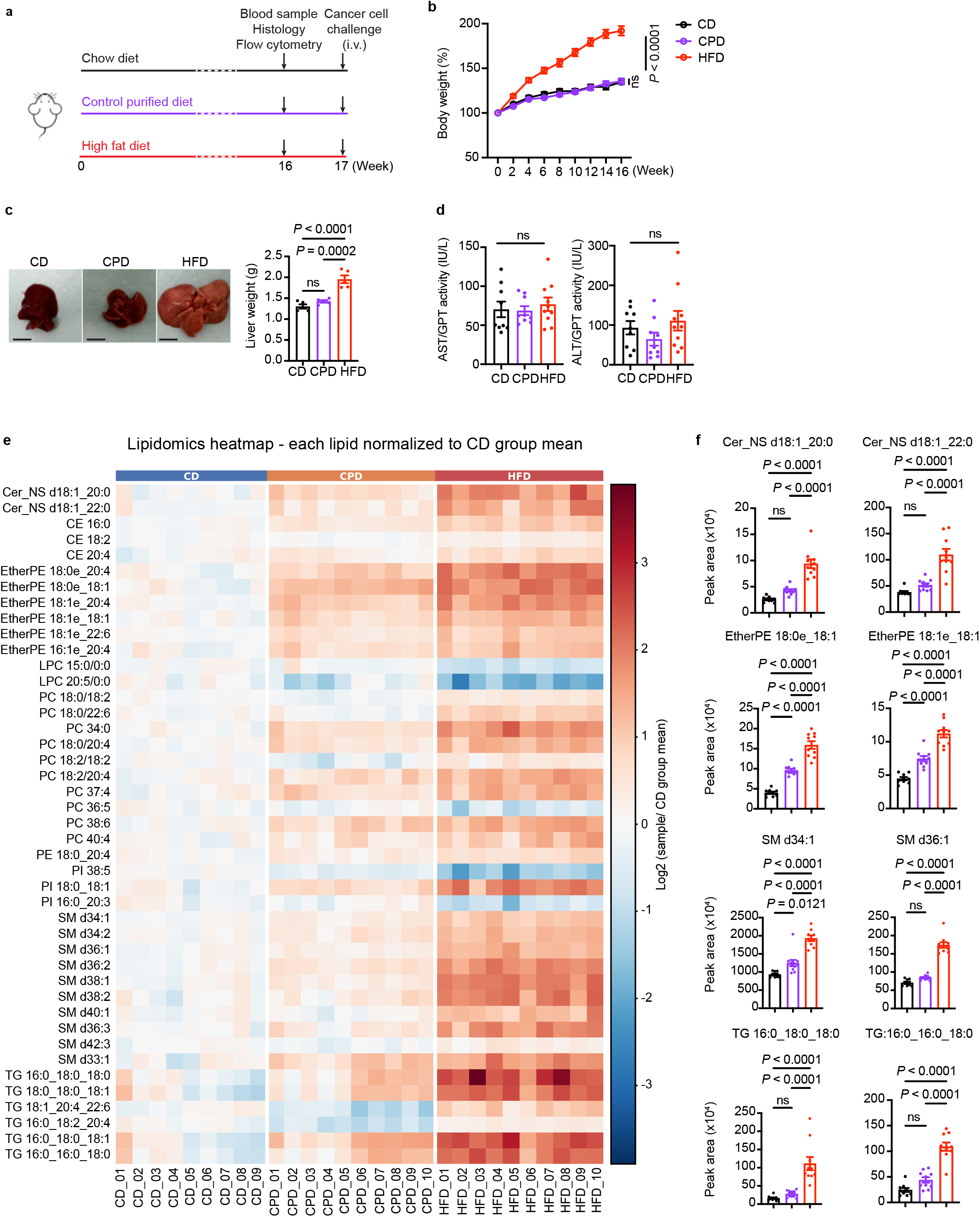
Effects of diets on body weight, liver weight and damage, and serum lipid and fatty acid composition. **a,** Schematic workflow illustrating dietary interventions, mouse phenotyping, and metastatic cancer cell challenge. **b**, Body weight over time in diet groups. Biweekly body weights of mice fed a control chow diet (CD; n = 10), a high-fat diet (HFD; 60% kcal fat; n = 10), or a control purified diet (CPD; calorie-matched with 10% kcal fat; n = 10) at the indicated timepoints. Mice were started on their respective diets at 8 weeks of age. *P* < 0.0001 for CD versus CPD and CD versus HFD; ns for CPD versus HFD. **c**, Representative liver images and liver weights of mice fed a CD (n = 5), CPD (n = 5), or HFD (n = 5) for 16 weeks. Liver weight quantification is shown as mean ± s.e.m. Scale bar: 10 mm. **d**, Circulating levels (serum) of aspartate aminotransferase (AST) and alanine aminotransferase (ALT) in mice maintained on a CD (n = 9), CPD (n = 10), or HFD (n = 10). AST and ALT level quantification is shown as mean ± s.e.m. **e**, **f**, Heatmap representing differential lipid species among the CD, CPD, and HFD groups, accompanied by quantitative analysis of selected lipid classes. Lipid concentration quantification is shown as mean ± s.e.m. *P* values were calculated using two-way ANOVA (**b**), Ordinary one-way ANOVA test (**c**-**d**, **f**), ns: not significant (*P* > 0.05).

**Extended Data Fig. 3.**
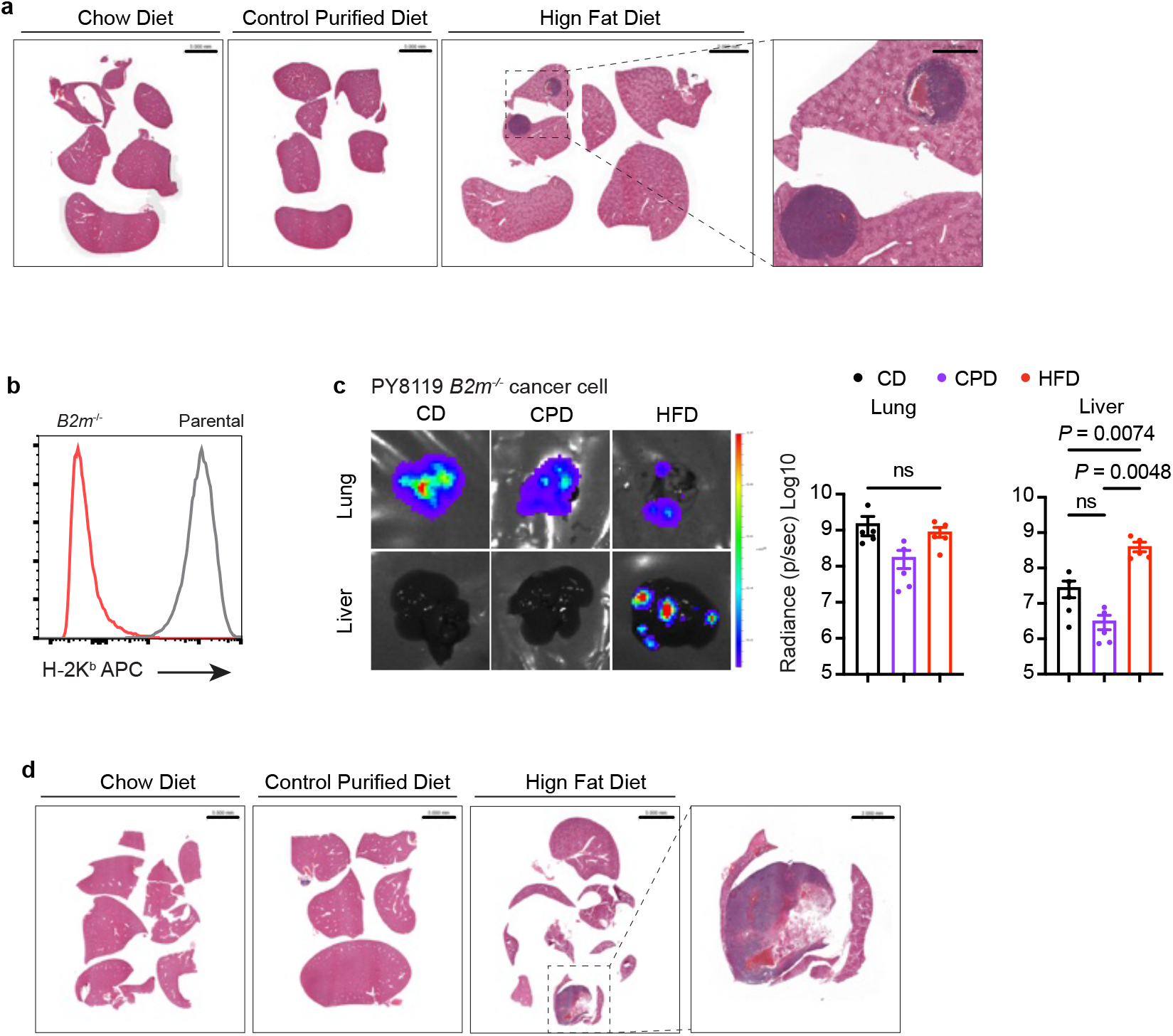
Breast cancer metastasis across dietary conditions. **a,** Representative hematoxylin and eosin (H&E) staining of liver sections from mice fed a control chow diet (CD), a high-fat diet (HFD, 60% kcal fat), or a control purified diet (CPD, calorie-matched with 10% kcal fat) and challenged with E0771 breast cancer cells. Scale bar: 5 mm. Magnified scale bar: 2 mm. **b**, Representative flow cytometry analysis of H-2K^b^ expression in parental and B2m-deficient (*B2m*^−/−^) PY8119 cancer cells. **c**, Representative bioluminescence images and radiance quantification of lung and liver colonization following intravenous administration of PY8119 *B2m*^−/−^ breast cancer cells in C57BL/6 mice fed a CD, CPD, or HFD (n = 5 per group). One representative image shown. Data are representative of at least three independent experiments. Radiance quantification is shown as mean ± s.e.m. **d**, Representative H&E staining of liver sections from mice fed a CD, CPD, or HFD following PY8119 *B2m*^−/−^ breast cancer cell challenge. *P* values were calculated using Ordinary one-way ANOVA test (**c**), ns: not significant (*P* > 0.05).

**Extended Data Fig. 4.**
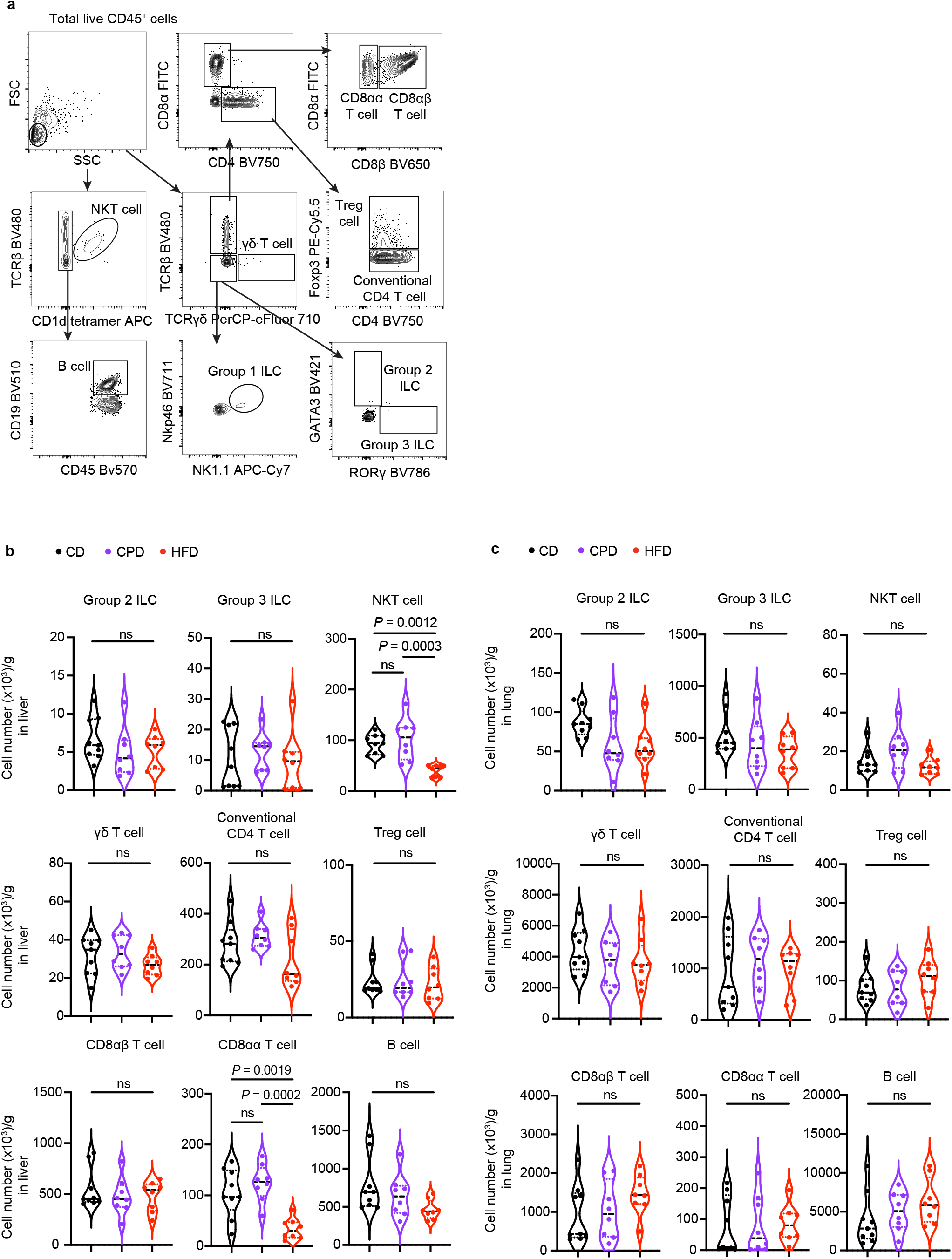
Effects of dietary feeding on innate and adaptive lymphocyte abundance in the liver and lung. **a**, Representative flow cytometry gating strategy used to define innate and adaptive lymphoid cell populations. **b**, **c**, Quantification of lymphocyte numbers in the livers and lungs of C57BL/6 mice fed a a control chow diet (CD, n = 9), a high-fat diet (HFD, 60% kcal fat; n = 8), or a control purified diet (CPD, calorie-matched with 10% kcal fat; n = 8). Cell numbers per gram of tissue are shown as mean ± s.e.m. Mice were analyzed without tumor challenge. *P* values were calculated using and Ordinary one-way ANOVA test (**b**, **c**). ns: not significant (*P* > 0.05).

**Extended Data Fig. 5.**
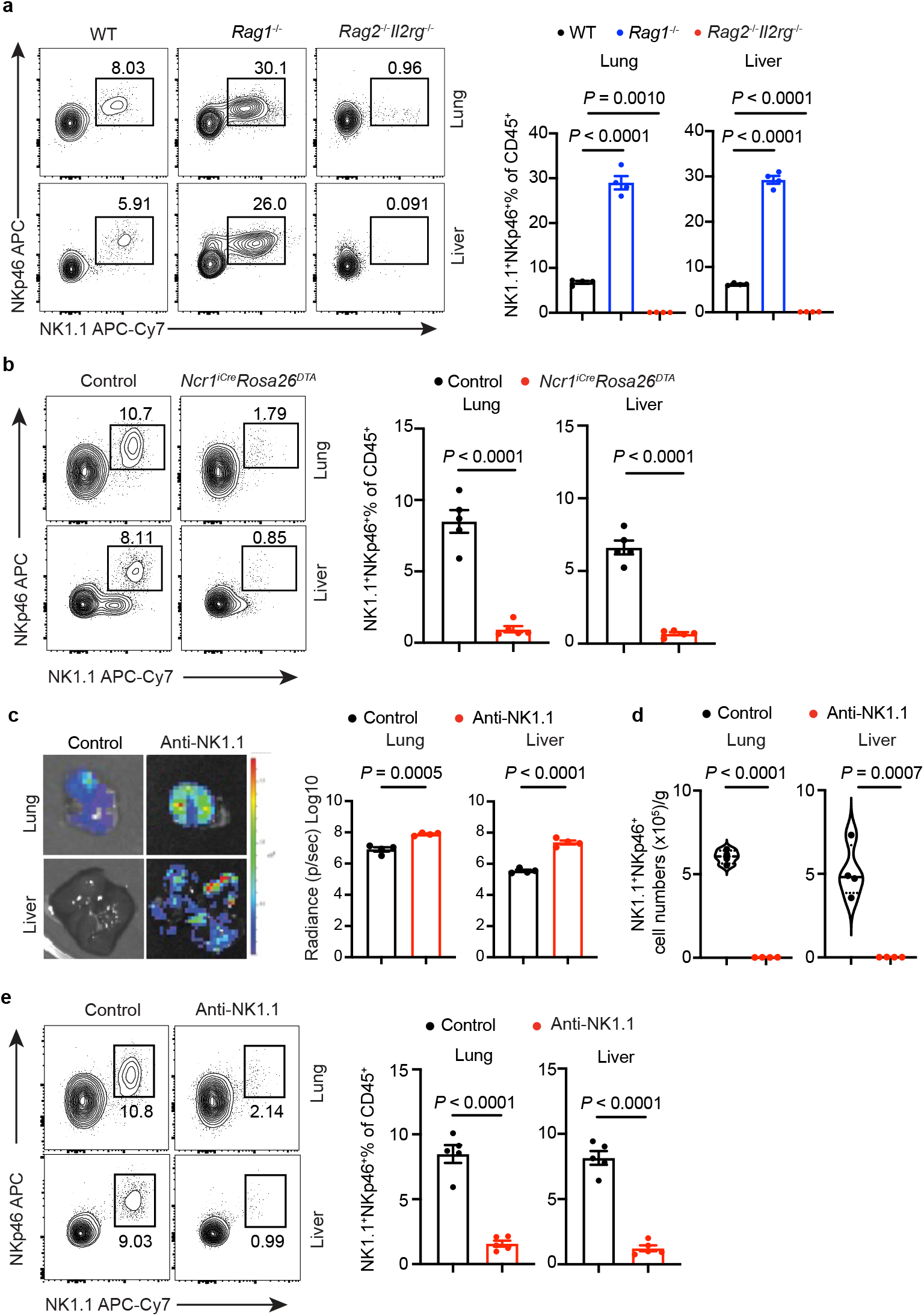
Group1 innate lymphoid cell depletion promotes breast cancer metastasis. **a**, **b**, **e**, Representative flow cytometry analysis and quantification of NK1.1^+^NKp46^+^ group 1 innate lymphoid cells in the lungs and livers of wild-type (WT), *Rag1^−/−^,* or *Rag2^−/−^Il2rg^−/−^*mice (**a**), *Ncr1^iCre^Rosa26^DTA^* and littermate control mice (**b**), and mice with or without anti-NK1.1 antibody (NK1.1 Ab) treatment (**e**) before E0771 cancer cell challenge (n = 5 per group). Proportions of NK1.1^+^NKp46^+^ cells among total CD45^+^ cells are shown as mean ± s.e.m. Mice were analyzed without tumor challenge. **c**, Representative bioluminescence images and radiance quantification of lung and liver colonization of cancer cells in C57BL/6 mice with or without anti-NK1.1 antibody treatment (n = 4 per group). Data are representative of at least three independent experiments; each cohort is n = 4 per group. One representative image shown. Radiance quantification is shown as mean ± s.e.m. **d**, Quantification of NK1.1^+^NKp46^+^ group 1 innate lymphoid cell numbers in the lungs and livers of C57BL/6 mice with or without anti-NK1.1 antibody treatment. Cell numbers per gram of tissue are shown as mean ± s.e.m. Mice were analyzed without tumor challenge. *P* values were calculated using Ordinary one-way ANOVA test (**a**), Welch’s *t*-test (**b**-**e**).

**Extended Data Fig. 6.**
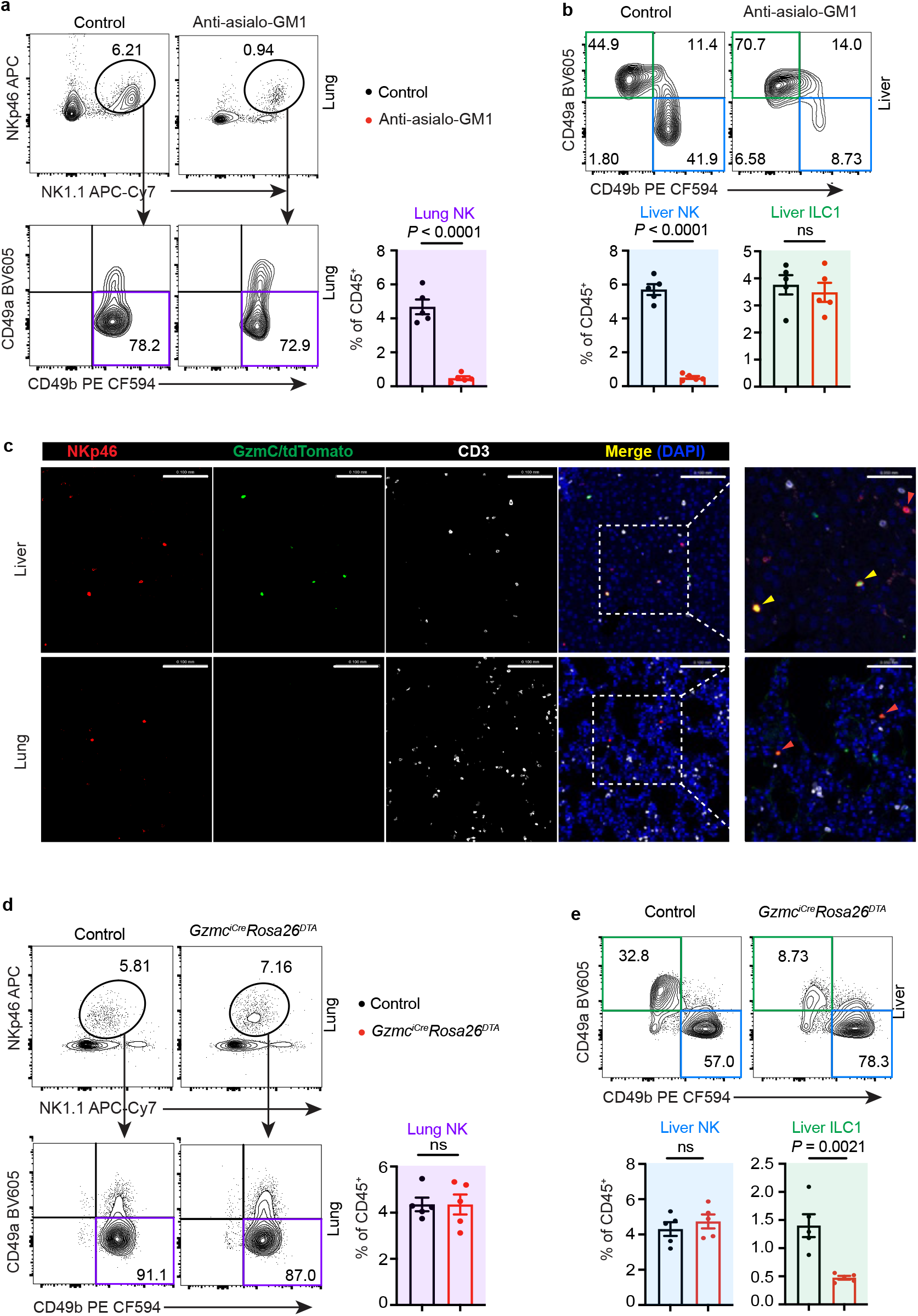
Anti-asialo-GM1 treatment and *Gzmc^iCre^Rosa26^DTA^* mice deplete NK cells and ILC1s, respectively. **a**, **b**, **d**, **e**, Representative flow cytometry analysis and quantification of NK1.1^+^NKp46^+^CD49a^+^CD49b^−^ ILC1s and NK1.1^+^NKp46^+^CD49a^−^CD49b^+^ conventional NK cells in the lungs and livers of C57BL/6 mice with or without anti-asialo-GM1 treatment (**a**, **b**) and *Gzmc^iCre^Rosa26^DTA^* and littermate control mice (**d**, **e**) (n = 5 per group). Proportions of ILC1s and NK cells among total CD45^+^ cells are shown as mean ± s.e.m. Mice were analyzed without tumor challenge. **c**, Representative images of GzmC^+^NKp46^+^CD3^−^ ILC1s (yellow arrowheads) and GzmC^−^ NKp46^+^CD3^−^ NK cells (red arrowheads) in the lung and liver of *Gzmc^iCre-tdtomato^*mice. Scale bar: 0.1 mm. Magnified scale bar: 0.05 mm. All mice were analyzed before tumor challenge. *P* values were calculated using Welch’s *t*-test. ns: not significant (*P* > 0.05).

**Extended Data Fig. 7.**
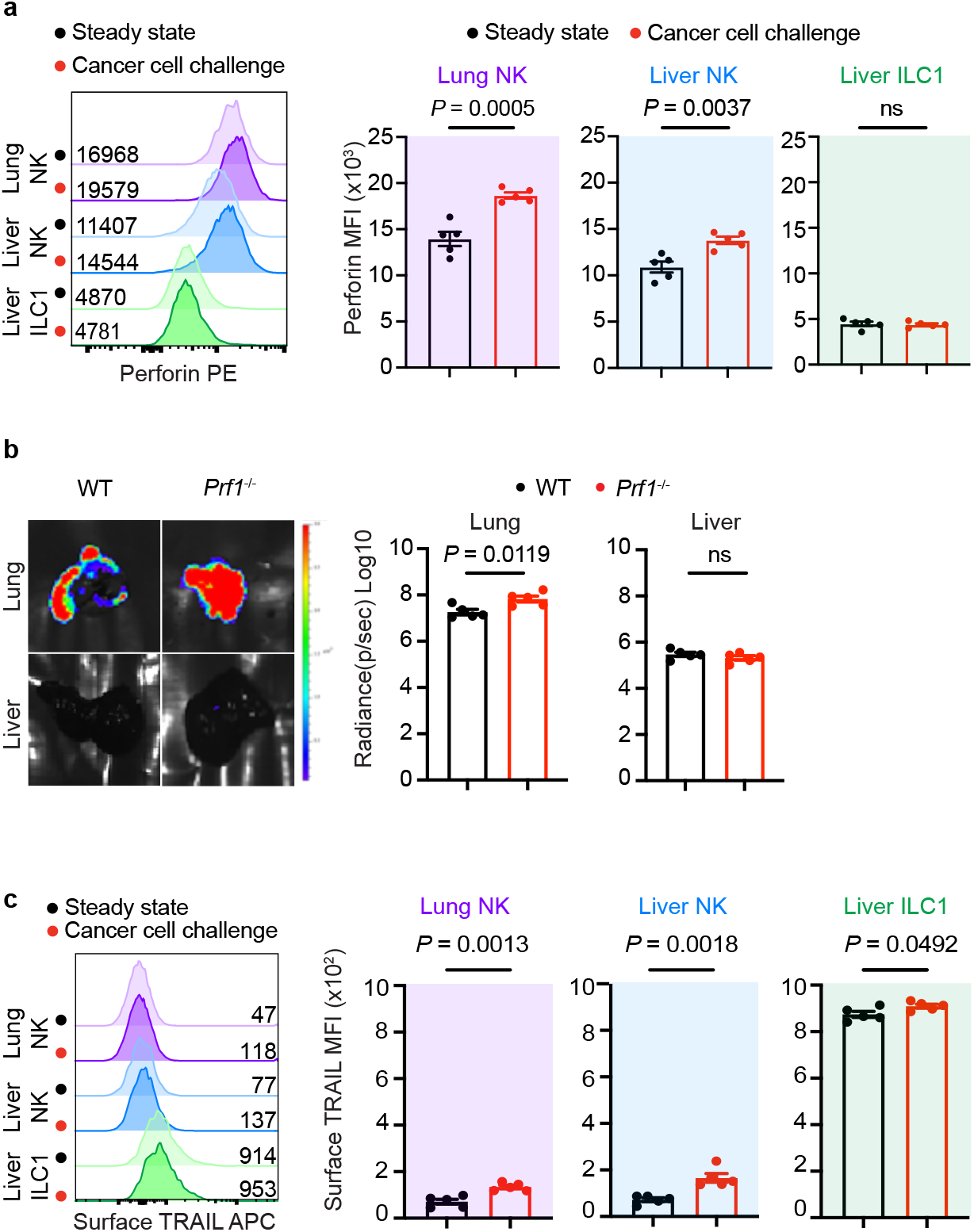
Expression and function of perforin and TRAIL in ILC1s and NK cells following cancer cell challenge. **a**, Representative flow cytometry analysis and quantification of perforin expression in ILC1s and NK cells in the lungs and livers of C57BL/6 mice with or without E0771 cancer cell challenge for 24 h. Perforin mean fluorescence intensity (MFI) is shown as mean ± s.e.m. **b**, Representative bioluminescence images and radiance quantification of lung and liver colonization of E0771 cancer cells in wild-type (WT) and *Prf1*^−/−^ mice. Data are representative of at least three independent experiments, and each cohort is n = 5 per group. One representative image shown. Radiance quantification is shown as mean ± s.e.m. **c**, Representative flow cytometry analysis and quantification of cell-surface TRAIL expression in ILC1s and NK cells in the lungs and livers of C57BL/6 mice with or without E0771 cancer cell challenge for 24 h. TRAIL MFI is shown as mean ± s.e.m. *P* values were calculated using Welch’s *t*-test. ns: not significant (*P* > 0.05).

**Extended Data Fig. 8.**
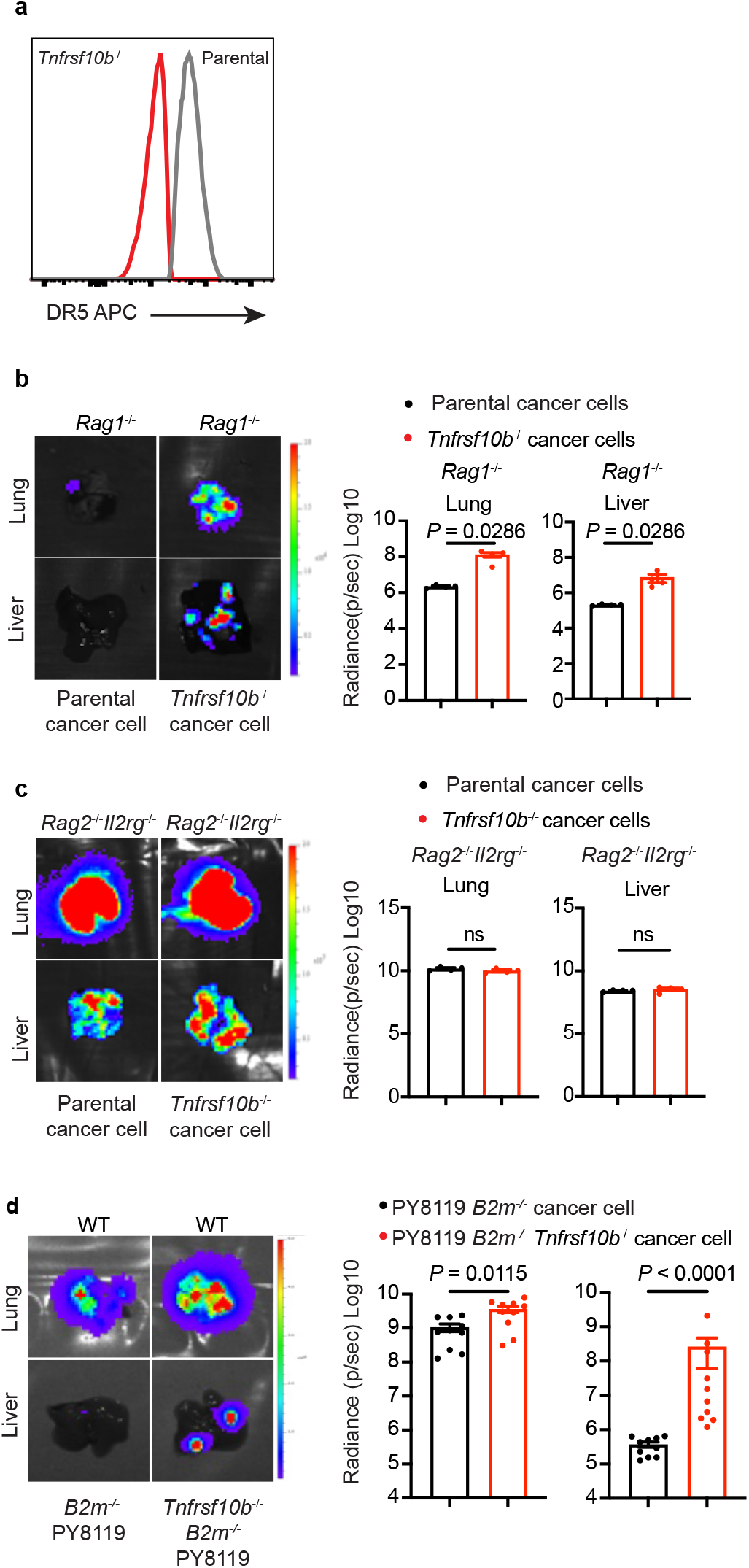
Accelerated metastatic growth of DR5-deficient breast cancer cells results from evasion of innate lymphocyte-mediated immunosurveillance. **a**, Representative flow cytometry analysis of DR5 expression in parental E0771 cancer cells, with DR5-deficient (*Tnfrsf10b*^−/−^) cells as a control. **b**, **c**, Representative bioluminescence images and radiance quantification of lung and liver colonization of parental and DR5-deficient (*Tnfrsf10b*^−/−^) E0771 cancer cells in *Rag1*^−/−^ (**b**, n = 4) and *Rag2^−/−^Il2rg^−/−^* (**c**, n = 4) mice. Data are representative of at least three independent experiments, each cohort is n = 4 per group. One representative image shown. Radiance quantification is shown as mean ± s.e.m. **d**, Representative bioluminescence images and radiance quantification of lung and liver colonization of parental (n = 10) and DR5-deficient (*Tnfrsf10b*^−/−^; n = 10) PY8119 *B2m*^−/−^ cancer cells. Data are representative of at least three independent experiments. Radiance quantification is shown as mean ± s.e.m. *P* values were calculated using Welch’s *t*-test. ns: not significant (*P* > 0.05).

**Extended Data Fig. 9.**
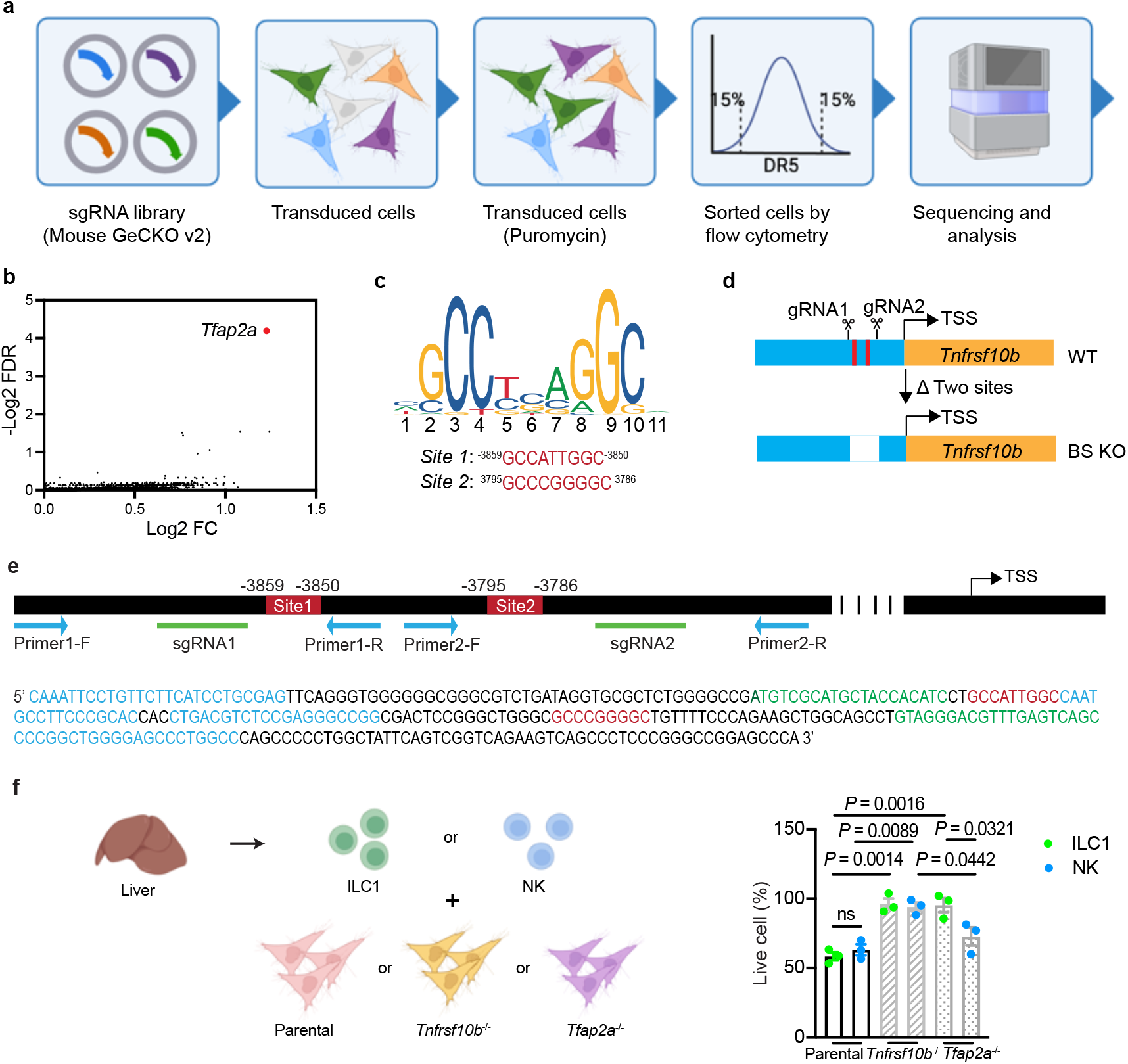
Regulation of DR5 expression in metastatic breast cancer cells and DR5-mediated cancer cell immunosurveillance by ILC1s and NK cells. **a**, Schematic overview of the CRISPR-Cas9 screening workflow for identifying DR5 regulators in E0771 cancer cells. **b**, Identification of AP-2α (encoded by *Tfap2a*) as a critical regulator of DR5 expression in E0771 cancer cells using a CRISPR-Cas9 screen. Volcano plot showing Log_2_ (fold change, FC) versus -Log_2_ (False Discovery Rate, FDR) of genes identified by Illumina sequencing analysis. Each point represents an individual gene. **c**, Top: Position-weight logo of the AP-2α binding motif (from JASPAR; https://jaspar.elixir.no/). Bottom: Predicted AP-2α binding sequences (Site 1 and Site 2) located upstream of the murine *Tnfrsf10b* transcription start site (TSS; position 0). **d**, **e**, Diagram (**d**) and sequence (**e**) illustrating the wild-type (WT) *Tnfrsf10b* gene locus and the generation of AP-2α binding site knockout (BS KO) cells using CRISPR-Cas9 genome editing. Predicted AP-2α binding sites are showing in red (**d**, **e**); sgRNA sequences in green; and genotyping primers used to identify BS KO cells in blue (**e**). **f**, (Left) Schematic diagram illustrating the experimental workflow for the in vitro functional assessment of hepatic group 1 innate lymphocytes. ILC1s and NK cells isolated from mouse liver were co-cultured with parental, *Tnfrsf10^−/−^*, or *Tfap2a^−/−^* luciferase-expressing E0771 cancer cells. (Right) Cytotoxicity was quantified as the percentage reduction in luciferase-mediated luminescence relative to control target cells cultured in the absence of effectors. The remaining viable target cell population (live cell percentage) is expressed as the mean ± s.e.m. *P* values were calculated using Ordinary one-way ANOVA test (**f**).

**Extended Data Fig. 10.**
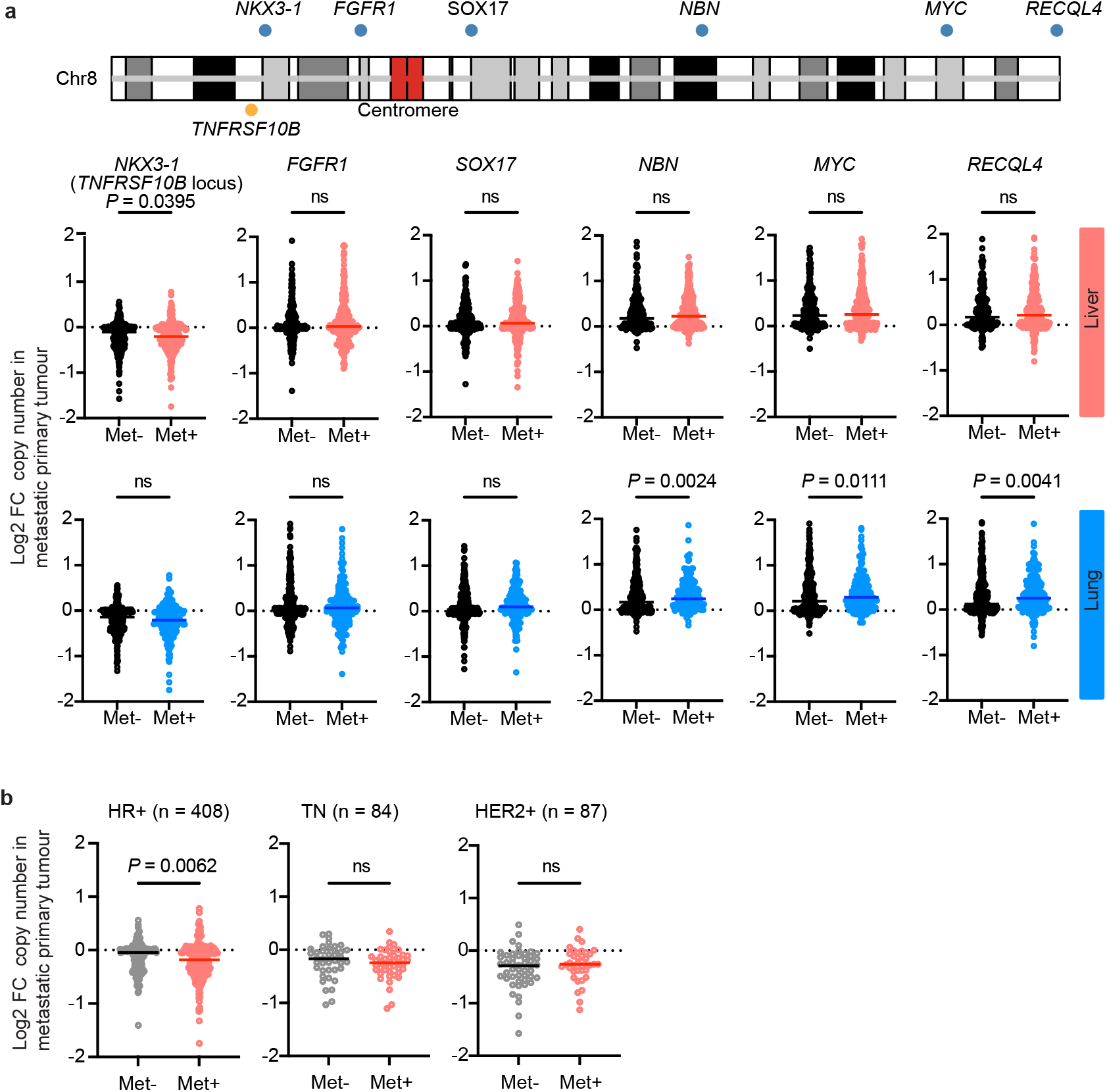
Gene locus copy number alterations in breast cancer patients. **a**, Selected chromosome 8 (Chr8) gene locus copy number alterations in breast cancer patients (MSK-IMPACT met_2021 cohort) with or without liver or lung metastases. The *TNFRSF10B* gene locus is closely linked to *NKX3-1*. Column plots show log_2_-transformed fold changes in copy number, with each dot representing an individual patient sample and the horizontal line indicating the median. **b**, Distribution of somatic copy number alterations at the *NKX3-1* gene locus across major clinical breast cancer molecular subtypes with or without liver metastasis: Triple-Negative (TN), Hormone Receptor-Positive (HR+), and HER2-Amplified (HER2+). Column plots show log_2_-transformed fold changes in copy number, with each dot representing an individual patient sample and the horizontal line indicating the median. *P* values were calculated using the Wilcoxon rank-sum test.

**Extended Data Fig. 11.**
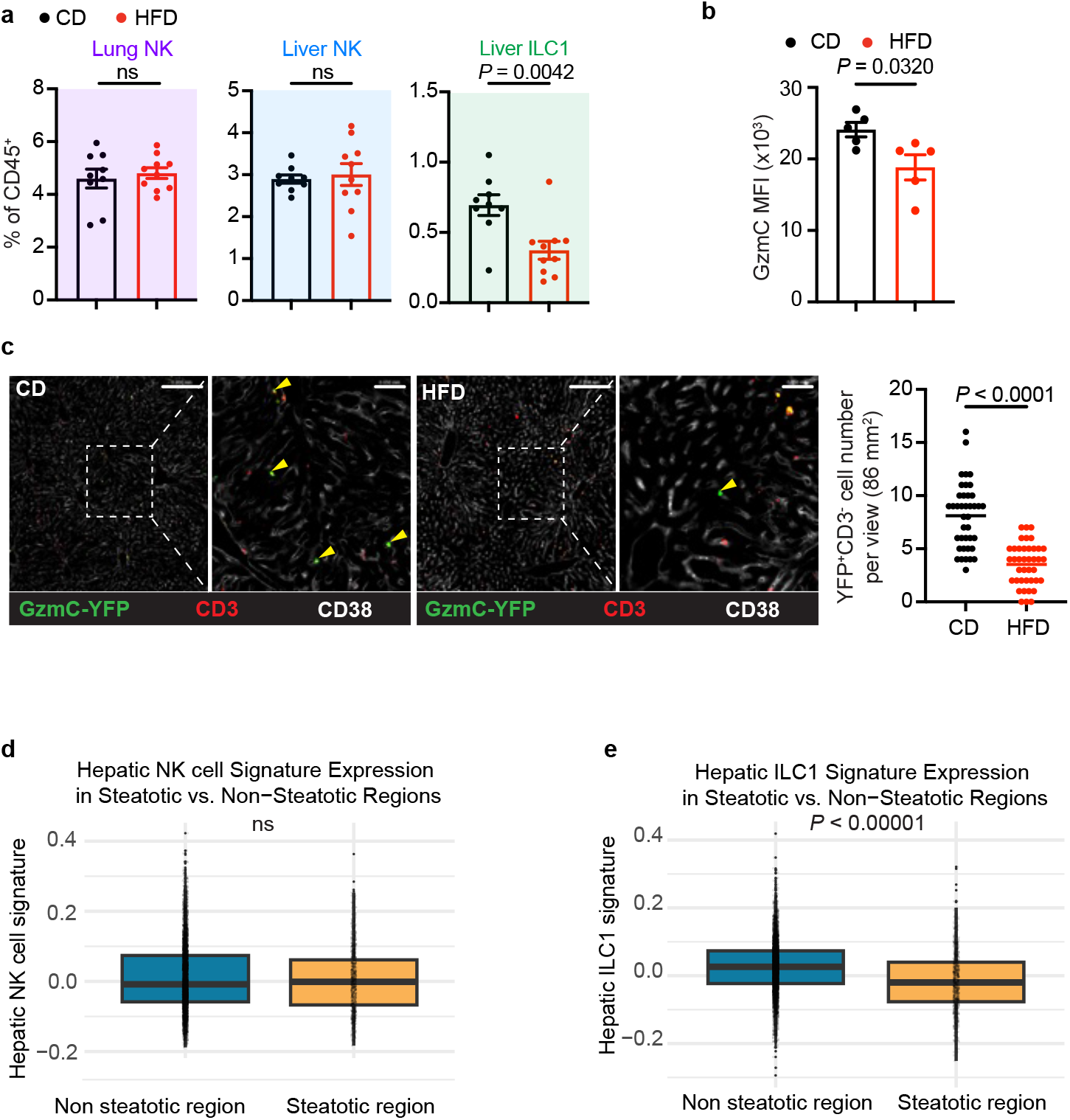
Hepatic ILC1s are selectively depleted in mice fed a high-fat diet and in human liver steatotic regions. **a**, Quantification of ILC1s and NK cells in the lungs and livers of C57BL/6 mice fed a control chow diet (CD, n = 9) or high-fat diet (HFD, 60% kcal fat; n = 10). Proportions of ILC1s and NK cells among total CD45^+^ cells are shown as mean ± s.e.m. Mice were analyzed without tumor challenge. **b**, Quantification of granzyme C (GzmC) expression in ILC1s from C57BL/6 mice fed a CD (n = 5) or HFD (n = 5). All mice were analyzed before tumour challenge. GzmC mean fluorescence intensity (MFI) is shown as mean ± s.e.m. Mice were analyzed without tumor challenge. **c**, Representative images and quantification of *Gzmc^iCre^* fate-mapped YFP^+^CD3^−^ ILC1s (arrowheads) in the livers of *Gzmc^iCre^Rosa26^YFP^* mice fed a CD (n = 2) or HFD (n = 2). A total of 40 fields were quantified per condition. YFP^+^CD3^−^ cell number per field are shown as mean ± s.e.m. Mice were analyzed without tumor challenge. Scale bar: 0.2 mm. Magnified scale bar: 0.05 mm. **d, e**, Expression of an NK cell gene signature (**d**) and ILC1 gene signature (**e**) in Visium regions annotated as steatotic versus non-steatotic, based on H&E review and unbiased clustering of spatial transcriptomic spots from human liver samples. *P* values were calculated using Welch’s *t*-test. ns: not significant (*P* > 0.05).

**Extended Data Fig. 12.**
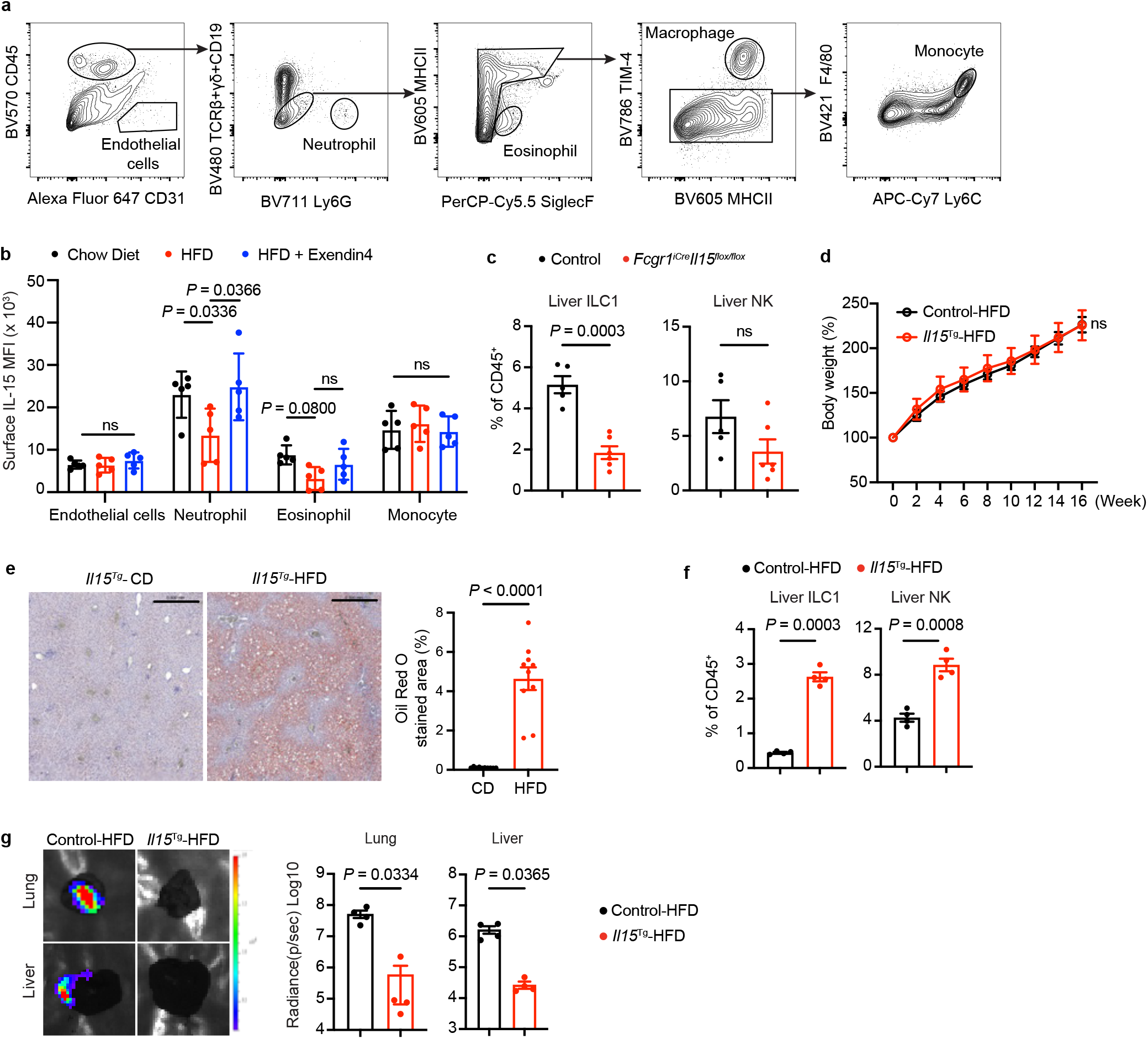
IL-15 is essential for the maintenance of liver-resident ILC1s and is reduced in the liver of high-fat diet-fed mice. **a**, Representative flow cytometry gating strategy used to define endothelial cell and myeloid populations in the liver. **b**, Quantification of cell-surface IL-15 expression in indicated populations from livers of mice fed a control chow diet (CD; n = 5), a high-fat diet (HFD, 60% kcal fat; n = 5), or an HFD with Exendin-4 treatment (n = 5). IL-15 mean fluorescence intensity (MFI) is shown as mean ± s.e.m. Mice were analyzed without tumor challenge. **c**, Quantification of ILC1s and NK cells in the livers of control (n = 5) and *Fcgr1^iCre^Il15^flox/flox^* (n = 6) mice. Proportions of ILC1s and NK cells among total CD45^+^ cells are shown as mean ± s.e.m. Mice were analyzed without tumor challenge. **d**, Body weight over time in control (n = 4) and *Il15^Tg^* (n = 4) mice fed an HFD. Biweekly body weights are shown at the indicated time points. Mice were started on an HFD at 8 weeks of age. **e**, Representative images of Oil Red O staining and hematoxylin and eosin (H&E) staining, along with Oil Red O quantification, of liver tissue from *Il15^Tg^* mice fed either a CD or an HFD. Oil Red O staining quantification is shown as mean ± s.e.m **f**, Quantification of ILC1s and NK cells in the livers of control (n = 4) and *Il15^Tg^* (n = 4) mice fed an HFD. Proportions of ILC1s and NK cells among total CD45^+^ cells are shown as mean ± s.e.m. Mice were analyzed without tumor challenge. **g**, Representative bioluminescence images and radiance quantification of lung and liver colonization of E0771 cancer cells in control (n = 4) and *Il15^Tg^*(n = 4) mice fed an HFD. One representative image shown. Radiance quantification is shown as mean ± s.e.m. *P* values were calculated using Welch’s *t*-test (**b-c, e-g**) and two-way ANOVA (**d**). ns: not significant (*P* > 0.05).

**Extended Data Fig. 13.**
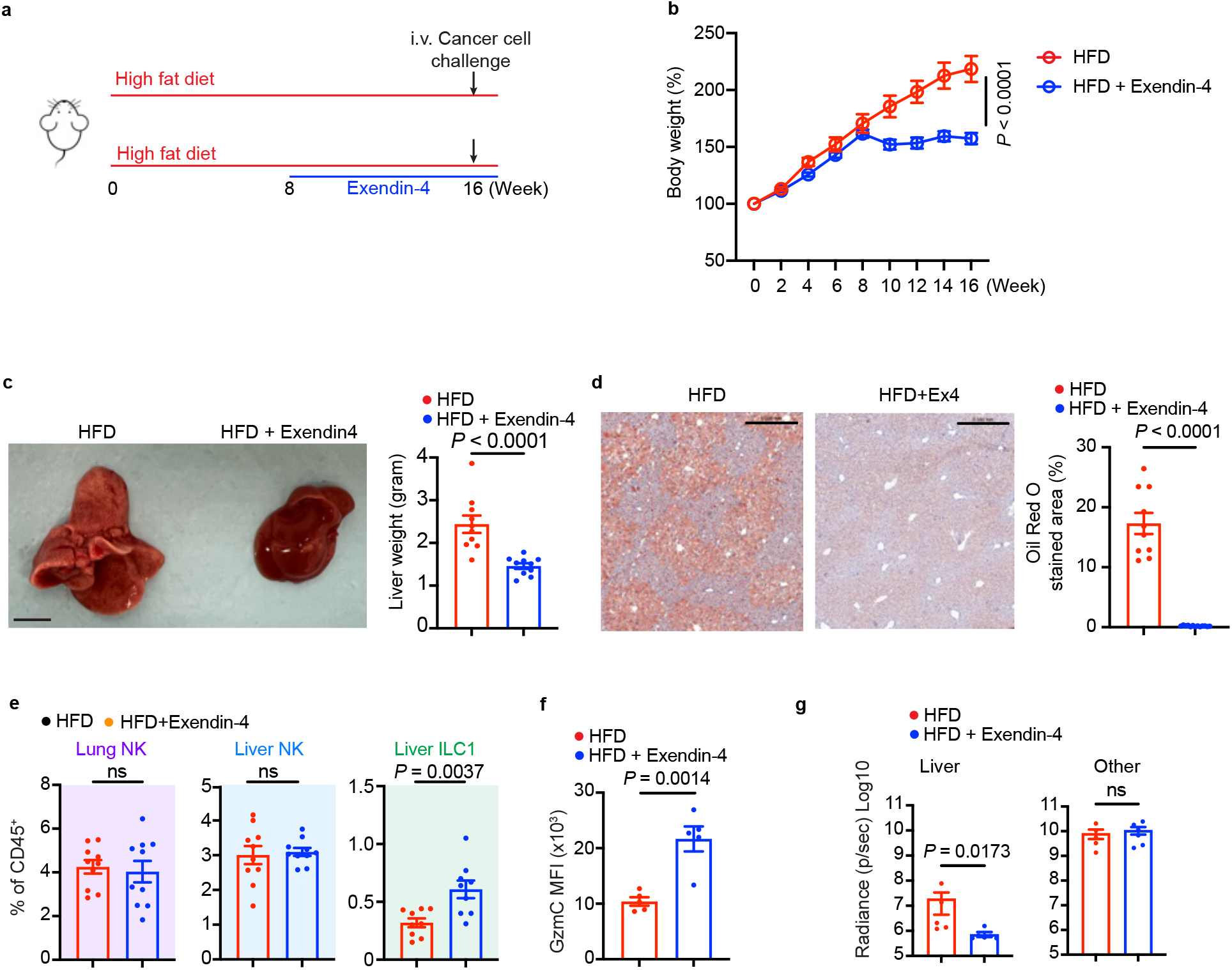
Exendin-4 treatment restores hepatic ILC1 populations in mice fed a high-fat diet. **a**, Schematic workflow illustrating the dietary interventions, Exendin-4 treatment, tumour challenge, and metastasis monitoring. **b**, Body weight over time in mice fed a high-fat diet (HFD, 60% kcal fat) with (n = 10) or without (n = 10) Exendin-4 treatment. Biweekly body weights are shown at the indicated time points. Mice were started on HFD at 8 weeks of age, and Exendin-4 was administered daily from weeks 8 to 16 of feeding. **c**, Representative liver images and liver weights of mice fed an HFD with (n = 10) or without (n = 10) Exendin-4 treatment. Liver weight quantification is shown as mean ± s.e.m. **d**, Representative images of Oil Red O staining and hematoxylin and eosin (H&E) staining, along with Oil Red O quantification, of liver tissue from mice fed an HFD with or without Exendin-4 treatment. Oil Red O staining quantification is shown as mean ± s.e.m **e**, Quantification of ILC1s and NK cells in the lungs and livers of C57BL/6 mice fed an HFD with (n = 10) or without (n = 10) Exendin-4 treatment. Proportions of ILC1s and NK cells among total CD45^+^ cells are shown as mean ± s.e.m. Mice were analyzed without tumor challenge. **f**, Quantification of granzyme C (GzmC) expression in ILC1s from C57BL/6 mice fed an HFD with (n = 5) or without (n = 5) Exendin-4 treatment. GzmC mean fluorescence intensity (MFI) is shown as mean ± s.e.m. **g**, Quantification of metastatic lesions detected in the liver and other sites in HFD with (n = 6) or without (n = 5) Exendin-4 treatment. Other sites evaluated include the lung, intestine, brain, and the abdominal region (including general abdominal tissue and abdominal white adipose tissue) *P* values were calculated using two-way ANOVA (**b**) and Welch’s *t*-test (**c**-**g**). ns: not significant (*P* > 0.05).

## Extended data table legend

**Extended Data Table 1 Lipidomics profiling of mouse serum across diet groups.**

Comprehensive serum lipidomics analysis from mice fed a control chow diet (CD), a high-fat diet (HFD, 60% kcal fat), or a control purified diet (CPD, calorie-matched with 10% kcal fat). The table details identified target lipid species that exhibited statistically significant differential abundance (upregulated or downregulated) in the HFD cohort compared to both the CD and CPD reference cohorts.

**Extended Data Table 2 DR5 regulators identified by a CRISPR/Cas9 screen.**

The table lists target gene identifiers and corresponding screen statistics, including log₂ fold change (LFC), false discovery rate (FDR), and –Log_2_(FDR), for each gene across the indicated conditions. LFC values represent the Log₂ fold change in sgRNA abundance relative to the control condition. FDR values were calculated using the Benjamini–Hochberg method.

